# Long deletion signatures in repetitive genomic regions track somatic evolution and enable sensitive detection of microsatellite instability

**DOI:** 10.1101/2024.10.03.616572

**Authors:** Qingli Guo, Jacob Househam, Eszter Lakatos, Salpie Nowinski, Ibrahim Al Bakir, Heather Grant, Vickna Balarajah, Christine S. Hughes, Luis Zapata, Hemant M Kocher, Andrea Sottoriva, Ann-Marie Baker, Ville Mustonen, Trevor A. Graham

## Abstract

Deficiency in the mismatch repair system (MMRd) causes microsatellite instability (MSI) in cancers and determines eligibility for immunotherapy. Here, we show that MMRd tumours harbour long-deletion signatures (≥2-5+ base pairs deleted in repetitive regions), which provide new insights into MSI evolution and enable sensitive MSI detection particularly in challenging clinical samples. Long deletions, accumulated through stepwise DNA slippage errors, are significantly more prevalent in metastatic MMRd tumours compared to primary tumours. Importantly, we show that long-deletion signatures harbour features that are distinct from background noise, making them robustly detectable even in shallow whole genome sequencing (sWGS, ∼0.1X coverage) of formalin-fixed samples. We constructed a machine learning classifier that uses these distinct features to detect Microsatellite Instability in LOw-quality (MILO) samples. MILO achieved 100% accuracy in detecting MSI in sWGS data with only 2%-15% tumour purity and demonstrated promise in identifying MMRd clones in precancerous intestinal lesions. We propose that MILO could be clinically used for the sensitive monitoring of MMRd cancer evolution from early to late stages, using minimal sequencing data from both archival and fresh-frozen samples with low tumour content.

**Significance:** Mutational signatures characterised by long deletions in repetitive genomic regions provide a sensitive route to detect and track MMRd clone evolution, even with low purity shallow whole genome sequencing data.

## Introduction

Mismatch repair deficiency (MMRd) is a well-recognised mutagenic process that has been found in cancers across more than 20 tissue types (1,2). MMRd cancers generally exhibit a more favourable prognosis and typically respond positively to immunotherapy (3,4). However, significant heterogeneity exists in clinical outcomes, including prognosis, response to chemotherapy, and sensitivity to immunotherapy (5,6). This heterogeneity highlights the urgent need for subtyping MMRd cancers and monitoring how MMRd tumours evolve over time and through therapy.

During DNA replication, the newly synthesised DNA strand can transiently dissociate from the template strand, causing misalignment and a phenomenon known as strand slippage error (7,8). Microsatellite (MS) regions, due to their repetitive DNA sequences, are particularly susceptible to these errors (8,9). This process causes the formation of small loops in DNA strands: if the loop forms on the new strand, it can cause an insertion, while on the template strand, it leads to a deletion (8).

The MMR machinery typically corrects post-replicative slippage errors; however, in cells with impaired MMR, this process is disrupted, resulting in microsatellite instability (MSI), characterised by a significantly elevated mutation rate of small insertions and deletions (indels) (10,11). Further, MMRd tumours accumulate single base substitutions (SBSs) in particular sequence contexts reflecting the intrinsic bias of the MMR system to identify and repair mismatched bases (12). Consequently, MMRd tumours show distinctive mutational signatures of both indels and SBSs (12,13).

Uncorrected slippage errors in MS loci create an ‘archive’, recording the evolutionary history of MMRd cancers (14,15). The burden of slippage events, termed ‘MSI intensity’ (usually measured as the proportion of mutated MS loci), is positively associated with immunotherapy response (6). Consequently, MSI characterisation may be useful to guide therapeutic decision-making (16). Current methods for MSI detection and characterisation struggle with challenging samples, particularly those with limited tumour cellularity and poor DNA quality, for example, samples from formalin-fixed paraffin-embedded (FFPE) materials (17,18).

Here, we explore the mutational signatures and evolution of MSI. We found that the accumulation of microsatellite slippage events serves as a surreptitious record of tumour evolutionary history and leads to distinct indel signatures, providing an exquisitely sensitive route for tracking MMRd tumours.

## Results

### Novel long-deletion signatures enriched in metastatic MMRd cancers

We characterised the spectrum of indel mutations in primary and metastatic MMRd cancers using deep sequencing of *n*=554 MMRd tumours from four publicly available datasets (EPICC (19,20): *n*=39; PCAWG (13,21): *n*=31; GEL (22,23): *n*=355; Hartwig (24): *n*=119), collectively comprising *n*=425 primary tumours and *n*=119 metastatic lesions. Samples included in the EPICC and GEL cohorts were derived from colorectal cancers (CRCs), whereas samples from the PCAWG and Hartwig cohorts were pan-cancer.

We derived the indel mutational spectrum for each cancer, represented as a vector of mutational frequencies across 83 mutation channels (13). These channels were determined based on the size of the indels and their surrounding sequence contexts. We then conducted a dimensionality reduction analysis using these indel profiles and observed a clear separation in the reduced-dimensional manifold between metastatic and primary MMRd cancers (Fig. 1a). By exploring the normalised indel profiles of CRCs in each cohort, we noted the prevalence of long deletions in the metastatic cohort that were absent from primary MMRd tumours (Supplementary Fig. 1).

**Fig. 1.**
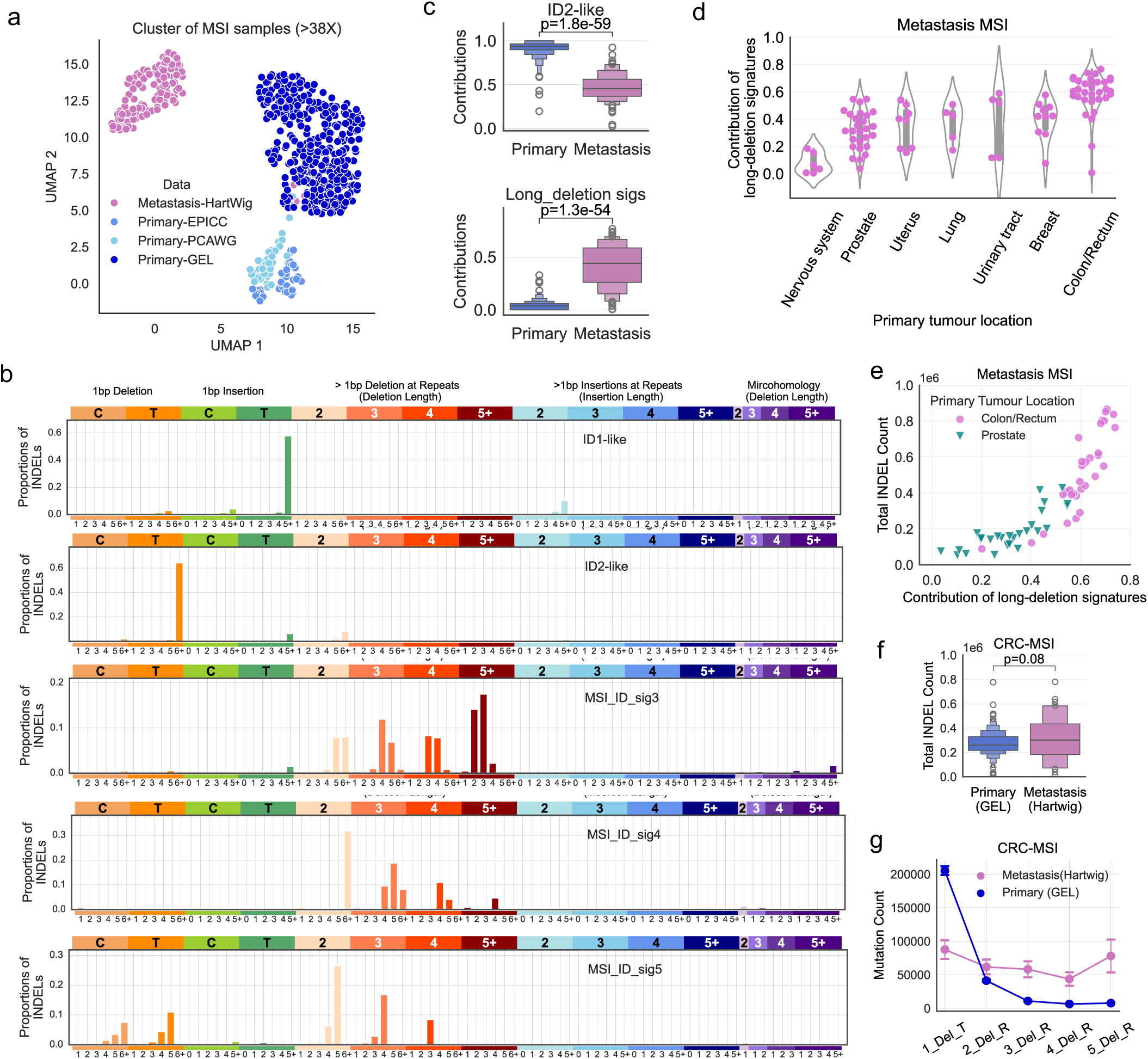
MSI indel signatures derived from deep-sequenced MMRd tumours. (**a**) Uniform Manifold Approximation and Projection (UMAP) cluster of combined MMRd cancers. We performed dimension reduction analysis on indel mutational profiles (83-channel) of 554 deep-sequenced MMRd cancers from four published data cohorts. (**b**) Inferred indel signatures from combined MMRd samples. (**c**) Contributions of dominant indel signatures in both primary and metastatic MMRd samples. The upper panel refers to ID2-like signature contributions and the lower panel refers to joint contributions of long-deletion signatures (MSI_ID_sig3-5). The statistical significance was assessed using a two-sided Mann–Whitney *U* test (the same for panel **f**). Data are presented using a Letter-Value plot and the black line corresponds to the median of the dataset and every further step splits the remaining data into two halves (the same for panel **f**). (**d**) Contributions of long-deletion signatures in metastatic MMRd cancers according to primary tumour locations. The width of the violin plot reflects the frequency of data points in each region, and the overlaid box plot in the centre indicates the interquartile range. (**e**) Correlation between long-deletion signature contributions and the total indel count in colorectal and prostate MMRd metastases. (**f**) Boxplot of total indel count of primary and metastatic CRC MMRd cancers. GEL cohort: 355 cases; Hartwig cohort: 34 cases. (**g**) The mean deletion count by length for primary and metastatic CRC MMRd samples, using the same sample set as shown in panel (**f**). Data are presented as mean values within each category +/- 95% confidence interval.

Next, we performed *de novo* mutational signature extraction and identified five indel signatures from the combined MMRd datasets (Fig. 1b; Supplementary Fig. 2; Methods). Two of the newly-derived signatures closely matched known MSI signatures (COSMIC v3.4) (13): ID1-like and ID2-like, with cosine similarities of 0.98 to signatures ID1 and ID2, respectively. The ID1-like signature features dominant 1 bp insertions of T/A in homopolymer (HP) regions, while the ID2-like signature features dominant 1 bp deletions of T/A in HP regions.

The other three signatures (MSI_ID_sig3, MSI_ID_sig4, and MSI_ID_sig5) are characterised by predominant features of longer deletions (ranging from 2 bp to more than 5 bp) at repetitive regions (microsatellites). Collectively, we will refer to these signatures as the ‘long-deletion signatures’. Among them, MSI_ID_sig3 and MSI_ID_sig5 are distinct from previously reported ID signatures (Supplementary Fig. 3) (13), while MSI_ID_sig4 shares some similarities with signature ID12 (cosine similarity of 0.87), which has an unknown aetiology. We noted that in MSI_ID_sig3, the mutational frequencies of longer deletions increase with deletion lengths from 2 to over 5 bp, whereas in MSI_ID_sig4 and MSI_ID_sig5, these frequencies decrease as deletion lengths increase.

We assessed the contributions of these newly identified indel signatures in our primary and metastatic MMRd tumour cohorts (Supplementary Fig. 4). We found that the average contribution of the long-deletion signatures was significantly higher in metastatic tumours (∼42%) than in primary tumours (∼4%) (Fig. 1c). In contrast, the ID2-like signature significantly decreased in relative contribution from ∼92% in primary to ∼47% in metastatic MMRd cancers (Fig. 1c). We observed that ID1-like, MSI_ID_sig4, and MSI_ID_sig5 signatures generally made smaller mutational contributions, but their activity was relatively increased in metastatic tumours compared to primary tumours (Supplementary Fig. 4).

There was a markedly greater variability in the relative contribution of the long-deletion signatures among metastatic cancer cohorts (standard deviation (SD) of 0.2) compared to primary MMRd tumours (SD of 0.04) (Fig. 1c). We found that the origin of the primary tumour was a crucial factor affecting the contributions of long-deletion signatures in metastatic MMRd lesions (Fig. 1d). MMRd metastatic samples originating from a colorectal primary exhibited the highest prevalence of long-deletion signatures, with a mean proportion of 58.4%, followed by breast-origin tumours and urinary tract tumours with mean proportions of 39.8% and 37.7%, respectively (Fig. 1d). In contrast, nervous system MMRd metastases showed the lowest proportion, with only 7%.

Across colorectal cancers (*n*=34) and prostate cancers (*n*=27), the two types of cancer with the greatest number of metastatic MMRd samples in the Hartwig cohort, there was a strong non-linear positive correlation between the contributions of the long-deletion signatures and total indel mutation counts (Spearman’s *ρ* = 0.61 for CRCs, *ρ* = 0.85 for prostate cancers) (Fig. 1e). Focusing on CRCs, we observed that the overall number of deletions showed a slight but non-significant increase in metastases compared to primaries (*p* = 0.08; Fig. 1f). However, stratifying the deletions by length (1 ∼ 5+ bp), showed a higher prevalence of 1 bp deletions in primaries vs metastases, while metastatic cancers displayed a substantial enrichment of long deletions (Fig. 1g; *p* ≤ 1.6 × 10⁻⁵), in agreement with the relative activities of MSI_ID_sig2 and the long-deletion signatures (Fig. 1c).

### Long deletions are likely accumulated through stepwise slippage events over time

Our analysis revealed a significant increase in the frequency of long deletions in metastatic MMRd tumours compared to primaries (Fig. 1c & g). Previous work in MMRd cell lines and mouse models has shown that long deletions occur through stepwise slippage errors (14,15). Here, we examined evidence of repeated slippages in five patients in the Hartwig cohort (four with prostate cancer and one with CRC) who had multiple longitudinally collected metastatic samples.

The frequency of motif deletions was inversely proportional to deletion size, with deletions longer than 20 bp being extremely rare (0.03%; Supplementary Table 1). Stratifying deletions by odd vs even length showed that 90% of odd-length indels and 85% of short even-length indels (≤10 bp) were composed of mono-nucleotide repeats (1 bp motifs) (Supplementary Fig. 5). The majority (63.7%) of the longer even-length indels (>10 bp) were formed by di-nucleotide repeats (2 bp motifs; Supplementary Fig. 5b). Therefore, most of the longer indels in advanced MMRd cancers consist of multiple repeats of smaller motifs (1 or 2 bp).

We examined the evidence for accumulation of repeated slippage events. For each patient, we searched for unique MS loci where slippage events were detected at both time points, referring to these as ‘shared mutated MS loci’ (Fig. 2a). The ‘slippage size’ refers to the number of repeat motif units differing between the two indels. Across the five longitudinally sampled patients, an average of 57% of the mutated MS loci were shared within the paired samples from each patient, with a wide range spanning from 19% to 90% (Fig. 2b). This variation may reflect differences in the divergence time between samples or the intra-tumour heterogeneity.

**Fig. 2.**
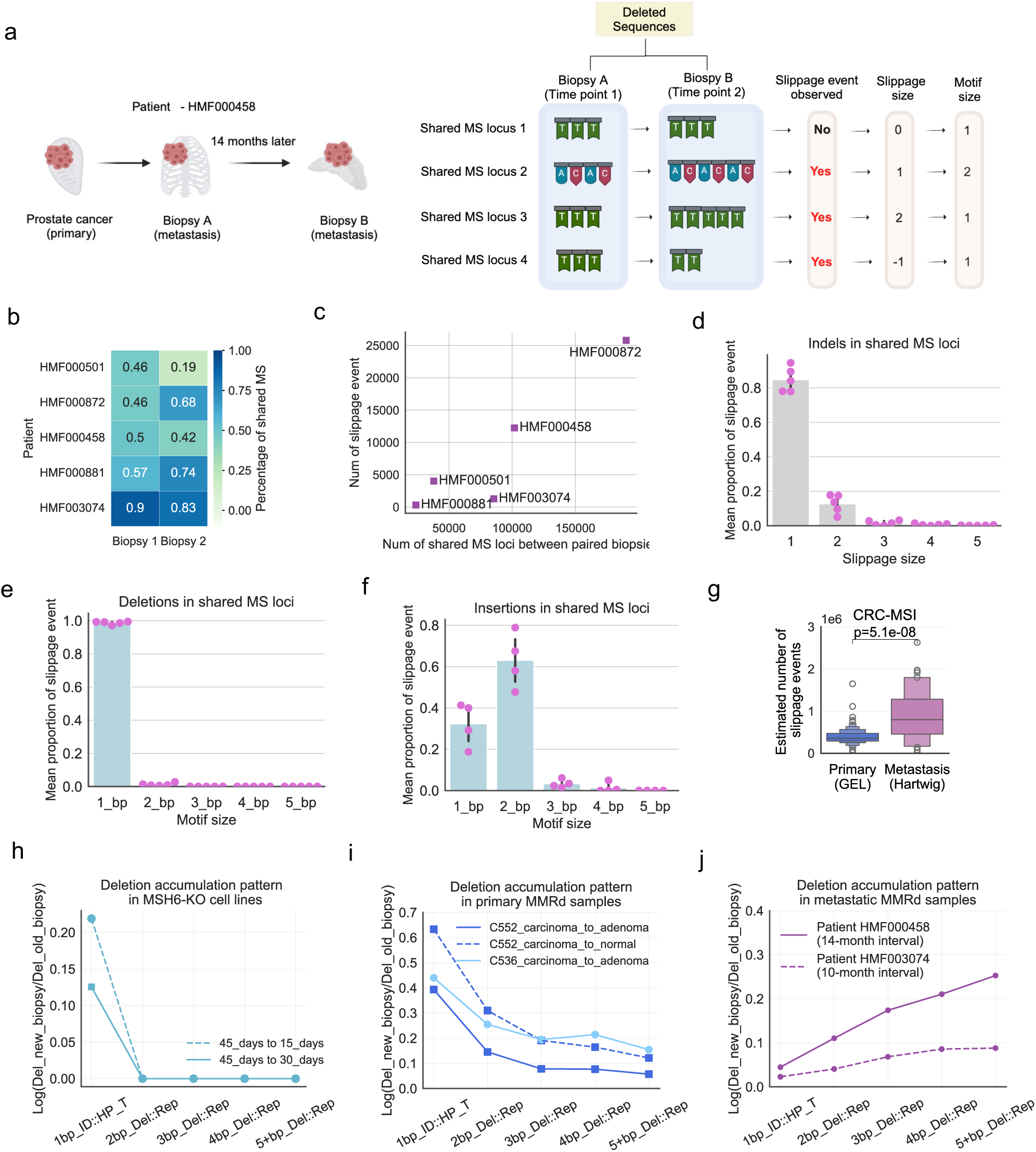
Stepwise accumulation of slippage errors leads to the formation of long deletions. (**a**) A schematic figure illustrating the definitions of ‘slippage event,’ ‘slippage size,’ and ‘motif size’ from deletions in shared MS loci. (**b**) Heatmap showing the proportion of shared MS loci in paired samples for five metastatic MMRd patients from the Hartwig cohort. (**c**) Correlation between the number of slippage events and the total number of shared mutated MS loci in the five metastatic MMRd patients. (**d**) The mean proportion of slippage events at different slippage sizes. Error bars representing the standard deviations. (**e-f**) The mean proportion of slippage events at different motif sizes for deletions (**e**) and for insertions (**f**). Standard deviations are shown as error bars. Data are presented for slippage events with observed numbers greater than 20. (**g**) Comparison of estimated total number of slippage events between primary and metastatic CRC MMRd cancers, using the same sample set as shown in Fig. 1f. Data are presented using a Letter-Value plot and the black line corresponds to the median of the dataset and every further step splits the remaining data into two halves. Statistical significance was assessed using a two-sided Mann-Whitney *U* test. (**h-j**) Deletion accumulation patterns by length: (**h**) in *MSH6* knockouts cell-lines; (**i**) in primary MMRd-CRC cancers; and (**j**) in metastatic MMRd cancer patients. A total of 8 samples were collected from *MSH6 deficient* stem cells. Our analysis uses the age of tumours following carcinoma > adenoma > normal-looking tissue.

Within the shared mutated MS loci, there was a substantial number of ‘new’ slippage events (mean of 8,752), where the later sample had a different number of motif repeats compared to the earlier sample, occurring in approximately 10% of shared MS loci across four out of five patients (Fig. 2c). New deletions (86%) were more common than new insertions (14%) on average.

We examined the slippage size across different motif lengths for all detected slippage events (Fig. 2d; Supplementary Fig. 6). Slippages of size one was the most common (with a mean of 85%) and the overall frequencies decreased exponentially with larger slippage sizes. The deletions primarily consisted of 1 bp motifs (98.7%; Fig. 2e). Additionally, 2 bp motifs expanded more frequently than 1 bp motifs, with a mean frequency of 63% and 32%, respectively, for sample pairs with more than 20 slippage events (Fig. 2f).

Notably, we found examples where an MS locus had undergone deletion of one motif at the first time point, but then later showed insertion at the same locus, and vice versa (slippage size < 0; Supplementary Fig. 6; Fig. 2a). It indicates that both insertions and deletions can occur within the same MS loci over time (a similar observation is reported in (25)), and the observed deletions and insertions from one biopsied sample are actually net changes after accounting for both types of events. Overall, due to the higher rate of deletions compared to insertions (12), we observed a prevalence of long deletions as the disease progresses. These observations likely result from repeated slippage events over time in clonal microsatellite mutations. We acknowledge that convergent evolution of different MMRd clones, although very rare, can also contribute to these findings.

Following the stepwise model, we expected that tumours with longer evolutionary time (since MMRd onset) and/or higher slippage rates would exhibit a greater number of total slippage events compared to those with more recent MMRd initiation or have slower slippage rates. We indeed observed a significantly higher number of slippage events in metastatic vs primary MMRd CRC tumours (Fig. 2g; Methods), whereas the total indel mutation burden alone was not significantly different (Fig. 1f). Overall, our results suggest that measurements incorporating the estimated number of slippage events may provide more informative insights into MSI evolution.

Next, we explored the indel accumulation pattern (calculated as log_10_ ratio of deletion numbers by size between later and earlier timepoints) in MMRd cell line and patient data. A stepwise slippage model predicts many 1 bp deletions of T/A in HP regions during the initial stage of MMRd onset and few deletions of 2+ bp (requiring slippage errors occurring at the same MS locus twice or more). Published data (12) where DNA was sequenced from a *MSH6* knockout (KO) cell line at three time points (15, 30, and 45 days post knockout), was consistent with the stepwise slippage model (Supplementary Fig. 7; Fig. 2h; Methods).

In our EPICC cohort of CRCs, we observed increased 1 bp deletion burden in MSI carcinomas compared to their precursor MSI adenomas, and a small increase in 2 bp deletions, with no noticeable difference in deletions >2 bp in two patients, C536 and C552 (Fig. 2i; Supplementary Fig. 8a-b). We found that histologically normal tissues from three MSI patients (C516, C548, and C552) exhibited a strong MSI signal (>10,000 1 bp T/A deletions in HP regions), albeit at lower levels than adenomas and carcinomas (Supplementary Fig. 8a, c & d). This suggests either contamination by tumour cells, or that MMRd can precede morphological changes, indicating a window for early detection of MMRd cancer clones.

In two metastatic cancer cases originating in the prostate (HMF000458 and HMF003047 from the Hartwig cohort), longitudinal biopsies were taken with an interval of 14 months and 10 months between samples, respectively. The later samples showed an increased burden of both short (1 bp) and long (2 ∼ 5+ bp) deletions (Fig. 2j; Supplementary Fig. 9a-b). The Hartwig cohort includes three other longitudinal sample pairs from MMRd cancers (without documented biopsy intervals) all of which showed a significant change in long deletions (Supplementary Fig. 9c-e).

### Higher long-deletion intensity correlates with better clinical outcomes in advanced MMRd cancers

The proportion of unstable MS loci has previously been shown to correlate with patient survival (26). We evaluated the potential of long deletions as a prognostic marker for patient outcomes (Fig. 3). We performed overall survival (OS) analysis using *n*=100 Hartwig samples (19 samples were omitted due to missing information). We calculated the ‘long-deletion intensity’ (Methods); this measurement considers both the prevalence and length of long deletions, with the latter providing an estimation for the number of slippage events.

**Fig. 3.**
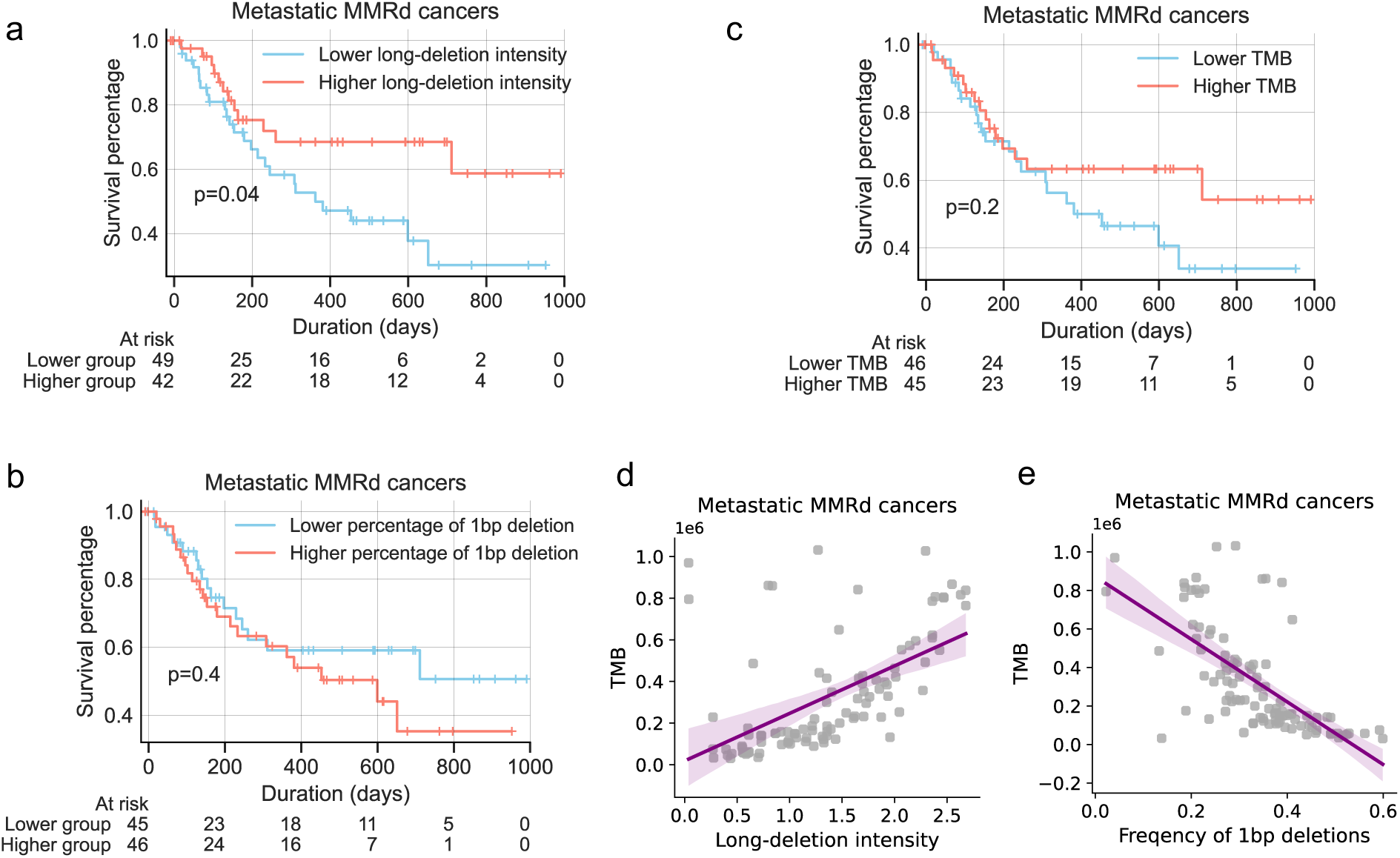
Higher long-deletion intensity is associated with better overall survival rate in metastatic MMRd cancers. (**a-c**) Kaplan-Meier overall survival curves of metastatic MMRd cancer patients stratified using several measurements: (**a**) long-deletion intensity, (**b**) total tumour mutation burden (TMB), and (**c**) frequency of 1 bp deletions. Long-deletion intensity is measured as the sum of (frequency of deletions × length of deletions) (Methods). TMB is calculated from the combined number of SBS and indels. The 1 bp deletion frequency refers to the normalised mutation frequency of 1 bp T/A deletions in HP regions (the dominant feature of the ID2 signature). The *p*-values were derived from the pairwise log rank test. (**d**) Correlation between long-deletion intensity and TMB. (**e**) Correlation between 1 bp deletion frequency and TMB. Purple lines in (**d-e**) represent the linear regression fits, with light purple bands showing the 95% confidence intervals.

We then assigned the samples into high (top 50%) and low (bottom 50%) long-deletion intensity subgroups (Fig. 3a). We found that Hartwig MMRd patients with high long-deletion intensity had significantly better overall survival compared to the low long-deletion intensity group (*p*=0.04; Fig 3a). In contrast, neither TMB nor the frequency of 1 bp deletions significantly stratified this cohort by overall survival (Fig. 3b-c). We observed that long deletions are enriched in high-TMB tumours whereas 1 bp deletions are progressively depleted as TMB increases (Fig. 3d-e), suggesting long deletions are more effective in predicting metastatic MMRd patient outcomes than a conventional marker such as TMB.

We also examined the association between long-deletion intensity and progression-free survival (PFS) using samples from 36 metastatic patients with documented treatment and response information. We found that when tumours were treated with chemotherapy, the majority (65%, 11 out of 17) experienced disease progression, regardless of their long-deletion intensity levels. However, when given therapies other than chemotherapy (19 cases, including nine who received immunotherapy), those with higher long-deletion intensity had longer PFS, although the difference was not statistically significant (Supplementary Fig. 10).

### Long-deletion signatures remain distinct MMRd signals even in low-quality samples

While primary MMRd cancers exhibit a lower prevalence of long-deletion signatures compared to metastatic MMRd cancers (Fig. 1c & g), they nonetheless harbour significantly more long deletions than MMRp tumours (Supplementary Fig. 11; *p* < 5.3 × 10⁻²³⁹). Although these differences are not visually discernible in indel signature profiles due to the dominance of ID2 (Supplementary Fig. 1), they suggest that long-deletion signatures may serve as ubiquitous markers for MMRd even in primary tumours.

Therefore, we explored the detectability of long deletions in shallow whole genome sequencing (sWGS) data from primary tumours. Firstly, we examined real and synthetic sWGS data from two CRC patients in the EPICC cohort: one with MMRd and one with MMRp. We down-sampled the reads from the deep-sequenced data (coverage of 45X) to generate synthetic sWGS (coverage of 0.1X and 0.01X) data (Methods). The EPICC cohort also contains genome data from the same tumours that were experimentally sequenced to 1X coverage. We applied a bioinformatics pipeline to call mutations in these sWGS samples (Methods), and considered variants as true somatic mutations (the ‘signal’) if they were also detected by the deep-sequenced data from the same cancer; otherwise, they were classified as ‘noise’ mutations.

We found that, at 1X coverage, the MMRd tumour, as a hypermutator phenotype, had significantly more true somatic mutations and a much higher signal-to-noise ratio (SNR) than MMRp tumour (Supplementary Fig. 12). Surprisingly, our synthetic ultra-shallow samples (coverage of 0.1X and 0.01X) exhibited similar SNRs compared to real sWGS samples with 1X coverage, and their signal mutation patterns were well preserved relative to the benchmark mutations (Supplementary Figs. 13-15). This suggested that MMRd signals were distinguishable even when highly diluted in sWGS data of primary MMRd tumours. We next explored if this finding held true in a larger cohort.

We examined a diverse large cohort of *n*=833 sWGS data obtained from FFPE as well as fresh frozen (FF) tissue biopsies (Fig. 4a). This cohort consisted of:

● *n*=265 FFPE biopsies from three independent studies: biopsies from patients with inflammatory bowel disease (UC-IBD; *n*=81) (27), duodenal cancers (*n*=120) and stage T1 colorectal cancers (*n*=64) (28). Among these samples, *n*=39 (14.7%) of samples were MMRd, as validated by immunohistochemistry for MMR proteins in prior studies. The mean coverage was approximately 0.1X for all cohorts.
● *n*=568 FF samples, of which 155 (27%) were MMRd from three previously published CRC cohorts: *n*=249 sWGS samples from individual glands of four CRC patients (Gland_lpWGS) (28), *n*=267 sWGS samples from the EPICC cohort (EPICC_lpWGS) (19,20), and *n*=52 down-sampled samples from deep-sequenced MSI cancers reported in our previous study (29), named ‘Mseq_ds_lpWGS’. The mean coverage of these three cohorts was 0.2X, 1.2X and 0.1X respectively.

**Fig. 4.**
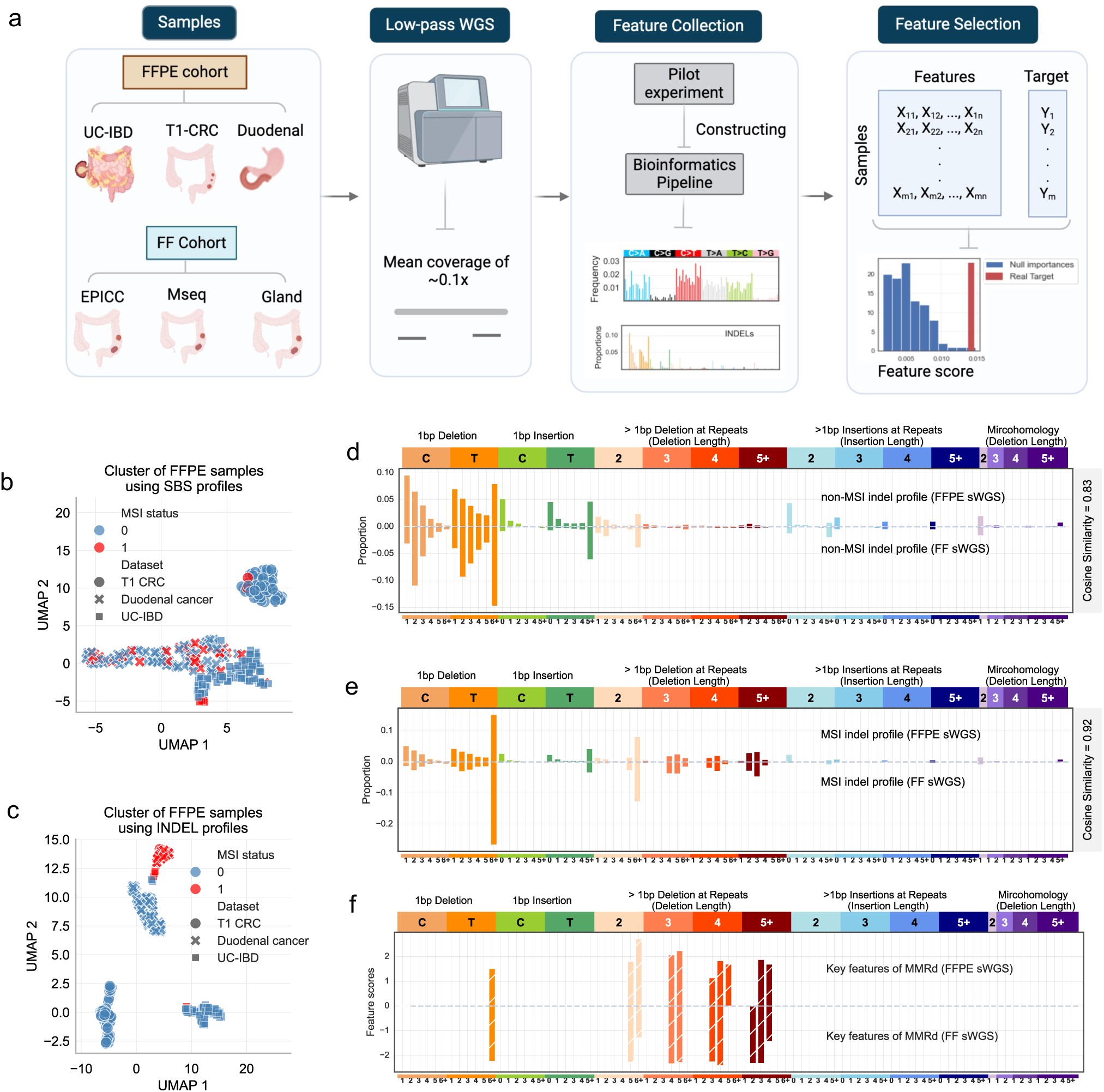
Identification of key signature features for MMR status in shallow whole genome sequence (sWGS) data. (**a**) A schematic representation of our data processing protocol. We collected 833 sWGS datasets from both FFPE and FF cohorts. We established a bioinformatics pipeline and applied it to all samples. We collected their SBS (96-channel) and indel (83-channel) mutational profiles, referred to as ‘features.’ Finally, we used a feature selection method to identify the most informative features of MMRd in sWGS samples. (**b-c**) Clustering of FFPE sWGS samples based on SBS (**b**) and indel (**c**) mutational profiles. We applied the dimension reduction method, Uniform Manifold Approximation and Projection (UMAP) using either their SBS profiles or indel profiles. The MSI status of ‘0’ refers to non-MSI samples (blue dots), and the MMR status of ‘1’ refers to MSI samples (red dots). (**d-e**) Average indel profiles of MMRp (**d**) and MMRd (**e**) samples derived from FFPE and FF sWGS cohorts. The *Y*-axis represents mutational frequency, while the *X*-axis denotes the 83 indel mutation channels. (**f**) Top ten most informative signature features for distinguishing MMR status in both FFPE and FF samples. The feature scores of the key signature features are represented by the bar heights. To distinguish these from the original indel mutation profiles in panels (**d**) and (**e**), white line strips were added to each bar.

We called SNVs and indels for all 833 sWGS samples using our established bioinformatics pipeline (Methods), and derived the mutational spectra for SBS (96-mutational channel) and indels (83-mutational channel). We then clustered FFPE samples (Fig. 4b-c) and FF samples (Supplementary Fig. 16) using SBS and indel mutation spectra separately. In both FFPE and FF cohorts, the indel profiles obtained from sWGS biopsies provided more insight into MMR status than the SBS profiles, probably due to the dominant background noise present in SBS mutation calls. The mean indel profiles for MMRd tumours were highly similar between FFPE and FF cohorts (cosine similarity of 0.92; Fig. 4e), whereas the similarity was lower for MMRp tumours (cosine similarity of 0.83; Fig. 4d). Batch effects on indel profiles were much more pronounced in MMRp samples in both the FF and FFPE sWGS cohorts (Supplementary Figs. 17-18).

We investigated which indel mutation features were most informative for MMRd status in sWGS data using a feature selection approach (30) (Methods). Among the top ten key features, nine were shared between FFPE and FF cohorts (Fig. 4e; Supplementary Fig. 19). These include the dominant ID2 signature peak and the long deletions (2 ∼ 5+ bp) in repetitive regions—which were also prominent in our long-deletion signatures derived from deep-sequenced MMRd tumours (Fig. 1b). These key features effectively distinguished MMRd from MMRp samples in both FF and FFPE sWGS data (Supplementary Fig. 20).

We noted that long-deletion signals were barely visible in indel profiles derived from deep-sequenced primary MMRd tumours (Supplementary Fig. 1a-c), but were readily detectable in indel profiles from our sWGS primary MMRd cases (Fig. 4e; Supplementary Figs. 17b & 18b). To investigate this discrepancy, we carefully examined each mutation calling step for 214 deep-sequenced EPICC samples. We found that long deletions were in fact called for all 32 MMRd tumours by the mutation caller, Mutect2 (31), but had been filtered out as they were labelled as ‘multiallelic’ without a ‘PASS’ tag (Supplementary Fig. 21). Long deletions were mostly enriched in multiallelic indels (Supplementary Fig. 21 c-d), providing further evidence for stepwise accumulation of short slippages leading to long deletions. Longer deletions had lower allele frequencies than shorter deletions, consistent with the idea that the repeat slippages occurred progressively (later) in tumour evolution. However, for sWGS samples, we included all detected mutations, which are typically covered by only a single read. As a result, multiallelism is extremely rarely detected; therefore, the ‘multiallelic’ flag was not used for filtering.

We compared the number of indel and SNVs that were filtered out in deeply sequenced MMRd and MMRp tumours (Supplementary Fig. 21a). We observed that indels were filtered out three times more frequently in MMRd cases compared to MMRp tumours, whereas the number of filtered-out SNVs was only slightly higher in MMRd tumours (Supplementary Fig. 21a-b). This strongly suggests over-filtering of indel mutations in MMRd tumours. When combining all indels detected in MMRd samples, the long-deletion signature features appeared at low frequencies, similar to what we observed in sWGS EPICC samples (Supplementary Fig. 21e-f).

Next, we sought to determine whether these long-deletion features enable sensitive detection of MMRd in low purity sWGS samples. We first examined 88 sWGS samples from six MMRd CRCs in the EPICC cohort (Fig. 5a). Of these, 47 were excluded in our original publication due to having ultra-low tumour cellularity and were not used in analysis shown in Fig. 4. Clustering analysis of the 88 indel signature profiles revealed a distinct separation between samples with low purity and those with higher tumour purity (Fig. 5a).

**Fig. 5.**
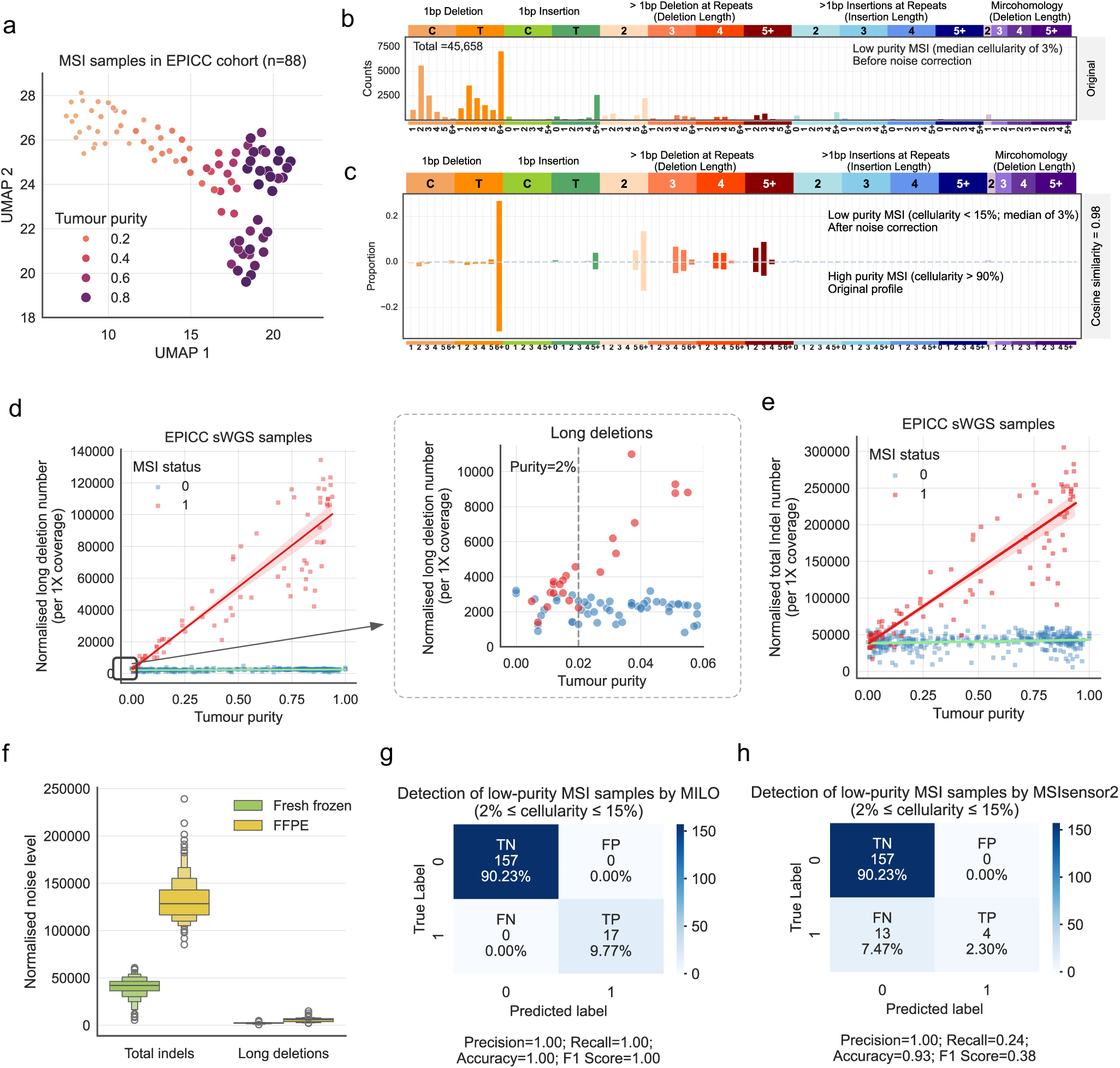
Long-deletion signature features are distinct MMRd signals and above background noise in low-quality samples. Low quality samples include low coverage, low-purity as well as archival samples. (**a**) Clustering 88 EPICC MMRd samples with various tumour purity. We employed the dimension reduction method, Uniform Manifold Approximation and Projection (UMAP), on their indel profiles and indicated tumour purity as the scatterplot diameter and continuous colour. (**b**) Averaged indel mutation profile of low-purity MMRd samples from the EPICC cohort. (**c**) Comparison of mean indel profiles between noise-corrected low-purity MMRd samples and high-purity MMRd samples. (**d**) Correlation between tumour purity and normalised long-deletion count in both MMRd and MMRp samples. The red and green lines show linear regression fits for MSI and non-MSI samples. The green line also indicates the average ‘noise’ (long deletions contributed by non-MSI cells). A zoomed-in view (indicated by a square box) highlights the MMRd detection limit at tumour purity of ∼2%. (**e**) Correlation between tumour purity and normalised total indel mutation count. (**f**) Comparison of the noise levels in MMRp samples between FFPE and FF cohorts. Data are presented using a Letter-Value plot and the black line corresponds to the median of the dataset and every further step splits the remaining data into two halves. (**g**-**h**) Confusion matrices showing the performance of MILO (**g**) and MSIsensor2 (**h**) on low-tumour purity samples. The matrices display true negatives (TN), true positives (TP), false positives (FP), and false negatives (FN).

The mean indel profiles of 32 low-purity MMRd samples (Fig. 5b; purity < 15%) demonstrated notably lower similarity to the mean high-purity MMRd profile (Fig. 5c; purity > 90%; *n*=9), with a cosine similarity of 0.74. Instead, they aligned closely with indel profiles in MMRp samples (cosine similarity of 0.99; top panel of Supplementary Fig. 18a), suggesting that the MMRd signal is diluted by the prevalent MMRp (stromal) cells in these low-purity biopsies. To unveil the masked MMRd signals in low-cellularity MMRd samples, we subtracted the MMRp ‘noise’ from the indel mutation profiles using the methodology from ref (32). This noise correction method treats the observed indel profile as a linear combination of signal (MMRd) and noise (MMRp) signatures, removing the noise indel signature to reveal the signal profile. The corrected profile closely resembled that of high-cellularity MMRd samples (cosine similarity of 0.98; Fig. 5c).

We then examined the correlation between indel mutation count and tumour cellularity to better understand the noise levels, particularly in low-purity samples (Fig. 5d-e; Supplementary Fig. 22). Here, we analysed a total of 474 sWGS samples from the EPICC cohort (consisting of 88 MMRd samples and 386 MMRp samples). This includes 267 samples presented in Fig. 4 and an additional 207 samples that were excluded from the original study due to ultra-low tumour purity. We noted that sequencing coverage could be a confounder: samples with higher sequencing coverage, regardless of their MMR status, exhibited a greater number of indel mutations (Supplementary Fig. 22). After coverage normalisation, we identified a strong correlation between tumour purity and both long deletion and total indel mutation counts in MMRd samples (Fig. 5d-e), whereas no such correlation was found with MMRd SNV mutations (Supplementary Fig. 23).

We observed that MMRp cells contribute a nearly constant number of indel mutations, which can be considered the background noise level in impure MMRd samples (Fig. 5d-e). Notably, the noise source for long deletions exhibited much less variance (SD of 568) compared to that of all indels (SD of 9,870). Additionally, formalin fixation introduced significantly less noise to long deletions compared to that introduced to other indels (Fig. 5f). These results indicate that long deletions stand out above all types of noise in these low-quality samples—such as unfiltered germline mutations, PCR and sequencing errors, and FFPE artefacts—making them ideal markers for detecting MMRd signals in low-quality samples. Based on these observations, we estimated that the detection limit for long-deletion signatures in low-purity MMRd samples is about 2% for FF samples and 5% for FFPE samples (Fig. 5d).

### MILO accurately predicts MMR status of low-purity sWGS samples

We consolidated these findings into a computational method termed ‘MILO’ (Detecting **M**icrosatellite **I**nstability in **LO**w-quality samples). MILO works by detecting the key signature features of MMRd identified in FF and FFPE cohorts, with most of these being long-deletion signature peaks (Fig. 4f). We simulated 20,000 FF and 20,000 FFPE samples by mixing MMRd (the ‘signal’) and MMRp (the ‘noise’) indel profiles together (Methods). For each cohort, half (10,000) of the samples were MMRd samples with varied tumour purities, and the other half were MMRp samples. We trained and tested three classifiers: Random Forest (RF), Supporting Vector Machine (SVM) and Naive Bayesian (NB), using the key features of the normalised indel profiles of the synthetic samples (Methods). Among them, RF demonstrated the best performance on the test datasets (Supplementary Fig. 24).

We then trained an RF classifier using synthetic samples generated from two of the three FF datasets and repeated the same process for FFPE cohorts (Methods). The trained classifier was tested on the third independent FF (*n*=267) or FFPE (*n*=120) dataset, achieving an accuracy of 99% for both cohorts (Supplementary Fig. 25). The RF classifiers detected one false-positive and one false-negative case in the third independent test datasets, with MSI probabilities of 0.74 and 0.61, respectively. Therefore, we categorised samples with an MSI probability between 0.5 to 0.75 as a low-confidence group in MILO (Supplementary Fig. 26).

To validate MILO’s calls, we examined an additional 174 EPICC sWGS samples (not used for training the classifiers), 17 of which were MMRd with low tumour purity ranging from 2-15%. MILO detected MMRd status with 100% accuracy on these samples (Fig. 5g), while in comparison, a widely used tool, MSIsensor2 (33), found only 4 out of 17 true MSI samples (24% recall rate; Fig. 5h).

### Colitis-associated MMRd cancer evolution: a case study of MILO application

People with inflammatory bowel disease (IBD) have an increased lifetime risk of CRC and are enrolled in surveillance programs to detect early signs of cancer, with multiple biopsies collected from the bowel, including from suspected pre-cancerous lesions (27). We collected 17 FFPE biopsies from three time points from a single IBD patient and performed sWGS at ∼0.1X on each biopsy (Fig. 6a).

**Fig. 6.**
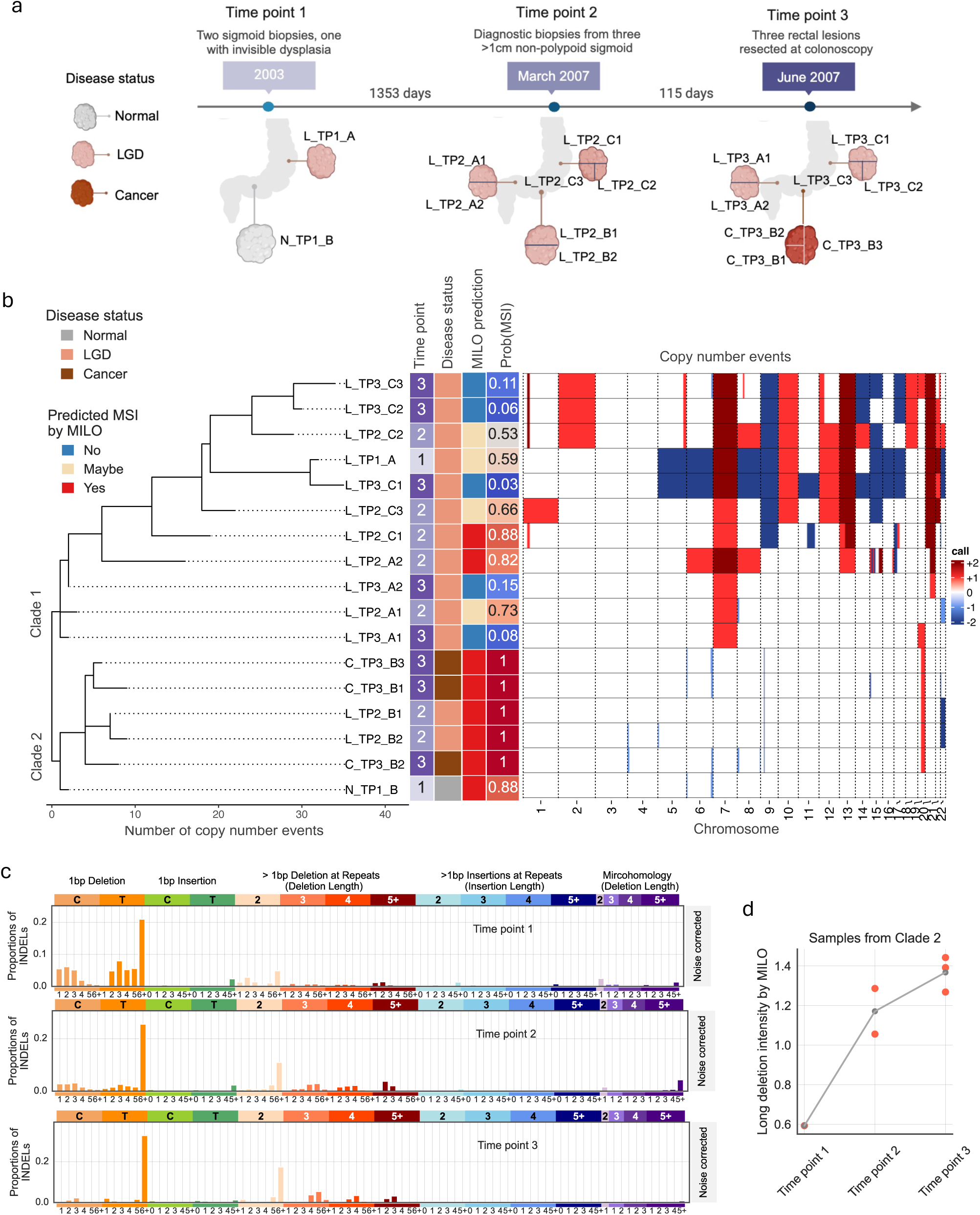
Early detection of MMRd clones in longitudinal IBD biopsies (FFPE sWGS) using MILO. (**a**) A schematic summarising the sample information. Biopsy IDs are annotated next to their approximate biopsy locations. (**b**) A phylogenetic tree constructed from the copy number profiles of 17 IBD biopsies, annotated with sample information and MILO prediction results. Copy number variations across 22 chromosomes are shown in the right panel. (**c**) Denoised indel profiles of clade 2 MMRd samples. The indel profiles at each time point were denoised and normalised (Methods). (d) Long-deletion intensities of clade 2 MMRd biopsies at three time points. Long-deletion intensity is predicted by MILO based on the noise-corrected indel profiles shown in panel (**c**) (Methods).

Aneuploidy is common in the colitic bowel (27), therefore we constructed a phylogenetic tree based on the copy number alteration (CNA) profiles from the 17 biopsies (Methods). This revealed two distinct clades (Fig. 6b), with clade 1 containing significantly more CNAs than clade 2. We then applied MILO to the indel profiles from each sample and detected MMRd signals in all six biopsies from clade 2. Clade 2 includes samples from all three time points, with a progressive increase in MSI intensity over time (as derived from their noise-corrected indel profiles; Fig. 6c-d; Methods). This suggests clonal expansion over time and through disease progression: at time point 1, the biopsy was classified as non-dysplastic, with the weakest MSI signal, indicating either that the MMRd clone was low cellularity or it accrued few indels; at time point 2, biopsies were classified as low grade dysplasia (LGD, a cancer precursor), and subsequently by time point 3, a CRC had developed.

In clade 1, MILO predicted that nearly half of biopsies (five out of eleven) were non-MSI. These non-MSI samples had extensive CNAs, which is consistent with the usual mutual exclusivity of CNAs and MSI (27,29). Two biopsies were assigned a high probability of being MSI (0.88 and 0.82), while four biopsies were categorised as having low-confidence MSI status (Fig. 6b). This potentially indicates that a second MMRd subclone had arisen in this patient but did not become clonally dominant.

## Discussion

In this study, we identified novel long-deletion signatures resulting from repeated slippage errors in microsatellite regions during MMRd cancer evolution. Although these long-deletion signatures occur at low frequencies (Fig. 4e), they collectively provide strong, robust, and unique MSI signals in pan-cancer MMRd tumours, especially in samples with low cellularity. This makes them ideal markers for detecting minimal residual disease (MRD) using cfDNA and for early screening of precancerous lesions using inexpensive sWGS data.

The prevalence of long deletions varies between individuals and across different tissues (Fig. 1). This may be due to factors such as tissue-specific mutation rates, cell division rates, and inter-tumour molecular differences, such as variations in MMR gene mutations (12,17,34–36). However, within an individual MMRd tumour, long-deletion signals generally increase with disease progression, offering a sensitive and reliable method for tracking tumour evolution.

We found that higher long-deletion intensity is associated with better clinical outcomes (Fig. 3a), likely because these samples are more antigenic. A similar result was observed by others, where a higher MSI intensity correlated with a better drug response rate, attributed to increased immune infiltrate in these samples (6). However, further investigation in a larger cohort is needed to establish the specific clinical utility of these quantitative measures.

Immunohistochemistry and PCR-based methods are gold standards for clinical MSI testing but have limitations with biopsies of low tumour cellularity (<30%) (37–40). Next-generation sequencing (NGS) provides a sensitive approach to MSI detection by analysing genomic features such as microsatellite length alterations, indel burden, and SBS profiles (12,33,41,42). However, these methods often require deep sequencing coverage (e.g., >20X) and high tumour purity, resulting in high costs and longer turnaround times, which limits their clinical practicality (43). MILO addresses these limitations by improving sensitivity in low-purity samples using cost-effective sWGS data, potentially making MSI calling by NGS more feasible for clinical use. Nevertheless, further studies are needed to evaluate its performance in low-purity non-CRC samples and its effectiveness in early cancer detection.

In conclusion, we identified novel indel signatures characterised by long deletions in repetitive regions, providing insights into their aetiology and demonstrating their utility in tracking MMRd cancer evolution. These signatures can be detected in challenging clinical samples using cost-effective data, making MILO a promising tool for early cancer detection and monitoring disease progression.

## Methods & Material

### Sample summary

#### Deep sequenced samples

The deep-sequenced MMRd cancers were sourced from four publicly available datasets: 1) Evolutionary Predictions in Colorectal Cancer (EPICC) cohort (*n*=39), including multi-region samples from six CRC patients with MSI (19,20); 2) Pan-Cancer Analysis of Whole Genomes (PCAWG) project by the International Cancer Genome Consortium (ICGC) (*n*=31), comprising primary MSI samples from individual patients with various cancer types (13,21); 3) Genomics England (GEL) 100,000 Genomes Project (100kGP; *n*=355), consisting of primary MSI cancers from patients with colorectal cancer (22,23); and 4) Hartwig Medical Foundation (*n*=119), encompassing metastatic MSI cancers from individual patients with different primary tumour types (24).

We also analysed indel mutations in a previous study observed from a knockout experiment of MMR gene, *MSH6*, (12). In their experiment, iPSC (induced pluripotent stem) cells were utilised, and the DNA was harvested for whole genome sequencing. We used *n*=8 samples from this dataset. Out of these samples, two were harvested after 15 and 30 days, while the remaining four samples were cultured for 45 days.

To collect indel mutation profiles, the VCF files from all deep sequenced samples were obtained from their original studies, and we employed SigProfilerMatrixGenerator (44) to collect their 83-channel indel profiles. Note that we used raw calling results of Mutect2 (31) from *n*=214 deep sequenced EPICC cohort to study the over-filtering issue of long deletions in Supplementary Fig. 21.

#### Shallow whole genome sequence (sWGS) samples

We analysed indel as well as SBS mutation profiles in sWGS samples from different sample cohorts, including both FFPE and FF samples. A total of 265 FFPE sWGS samples were collected from three independent studies. It includes: 1) *n*=81 samples from patients with inflammatory bowel disease (IBD) (27), encompassing 4 (∼5%) MSI samples. This study employed multi-region samples from 19 patients (27). The samples consisted of both non-cancerous colon tissues (low or high-grade dysplasia) and cancer tissues; 2) *n*=120 small bowel cancers in the duodenum with 32 (∼34%) MSI cancers. Each sample was obtained from an individual patient. Detailed information regarding this data cohort is described in a separate study (the manuscript is under preparation); and 3) *n*=64 from 34 CRC patients at the T1 stage, with 2 (∼3%) MSI cancers (28). The determination of MSI status was described in the respective studies.

We then collected 568 FF sWGS samples from three independent cohorts. This encompassed *n*=249 samples derived from individual glands in four CRC patients (two MSI and two MSS) with a mean coverage of 0.2X, referred to as the Gland_lpWGS dataset (28). The EPICC cohort contributed *n*=267 multi-region sWGS samples (mean coverage of 1.2X) from 67 CRC patients (19,20), encoded as the ‘EPICC_lpWGS’ dataset. Furthermore, *n*=52 synthetic low-pass data derived from 11 CRC patients (Mseq_ds_lpWGS) were included (29). The MSI status of each sample was annotated in their respective original studies. There are 101 out of 249 (∼41%), 41 out of 267 (∼15%) and 13 out of 52 (∼25%) MSI samples in Gland_lpWGS, EPICC_lpWGS and Mseq_ds_lpWGS, respectively.

Synthetic low-pass samples were generated for the Mseq_ds_lpWGS cohort and for a few samples used in the pilot study described in Fig. 4a using Samtools (45). Our target depth for the low-pass data was approximately 0.1X. To achieve this, we calculated the percentage of remaining reads based on the original depth, and it was then used in Samtools to randomly select reads from the deep-sequenced BAM files.

#### Low-purity sWGS samples

We analysed an additional 207 samples from the EPICC study that were filtered out in the original studies (19,20) due to their ultra-low tumour content, and 47 of 207 were MSI. The method of sample collection and processing is described in the original studies (19,20).

#### Longitudinal IBD samples

We also conducted a case study using 17 longitudinal samples from an IBD patient. FFPE human tissue samples, with corresponding anonymised clinical data, were obtained with ethical approval from St. Mark’s Hospital (NHS REC reference 18/LO/2051) from patients with extensive ulcerative colitis (Montreal Classification E3). Detailed information regarding the biopsy method, pathology classification and DNA library preparation protocol are summarised in this separate study (46). For this patient, multiple tissue biopsies were obtained at various time points using colonoscopy, targeting the neoplastic regions as determined by their attending physicians. Furthermore, copy number calling and phylogenetic tree reconstruction of these samples are also described in this separate study (46).

### Generating mutation profiles and signature analysis

#### Mutation calling for sWGS samples

We conducted a pilot study to determine the suitable bioinformatics pipeline for maximising true somatic mutations in sWGS data, which is then applied on all sWGS samples analysed in this study. Firstly, the BAM files for all sWGS samples were obtained. Most BAM files were aligned to the hg19 genome assembly using BWA-MEM (47), except for the EPICC cohort, which was aligned to the GRCh38 reference genome. To ensure data integrity, we verified that all BAM files were sorted by genome coordinate using SortSam (Picard), and duplicate reads were identified and marked using GATK’s MarkDuplicates (31). Additionally, we utilised CollectWgsMetrics to calculate the mean coverage for each processed BAM file. Subsequently, Octopus (48) was applied to all samples to generate a raw mutation list, providing the corresponding reference genome for each sample. We performed a filtering step on the generated VCF to exclude germline SNPs using ANNOVAR (49).

#### Mutational profiles

We employed SigProfilerMatrixGenerator (44) to collect their 96-channel SBS profiles and/or 83-channel indel profiles from the mutation list of all samples in this study. The mutation channels for both types of mutations are described and used in COSMIC v3.4 (https://cancer.sanger.ac.uk/signatures/id/). In this paper, we also projected deep-seq and sWGS samples on the first two principal components generated by the Uniform Manifold Approximation and Projection (UMAP) method using their indel or SBS mutation profiles.

#### Potential over-filtering of indels for deep sequenced samples

We went through the annotated raw mutation lists generated by Mutect2 of *n*=214 EPICC samples. Failed mutations are those without a ‘PASS’ tag. The final passed mutations are those reported in original EPICC studies. Further, the failed indel mutations were classified to two subgroups (‘multiallelic’ or ‘single allele’) based on whether there are more than two alternative alleles detected in the same MS loci. We then derived the mutation profiles from each subgroup for further analysis.

#### Mutational signature analysis

To infer indel signatures from deeply sequenced MMRd samples, we used a *de novo* signature analysis method similar to that described in (50). Briefly, we used two metrics to determine the number of active signatures: 1) cosine dissimilarity (1 - cosine similarity) between reconstructed and original mutation profiles, and 2) cosine similarities between all pairs of derived signatures to assess potential underfitting and overfitting scenarios. We applied the NMF deconvolution algorithm, implemented in the ‘sklearn’ package in Python, on the indel profiles of 554 deep-sequenced MMRd cancers using varying numbers of signatures (from 2 to 9). The above two metrics were derived from the solutions provided by the NMF algorithm for each model. To determine the actual number of active signatures, we choose the solution with lower reconstruction dissimilarity as well as a relatively lower cosine similarity between the inferred signatures. The inferred signature activities were further obtained using the refitting function implemented in our previously developed tool, FFPEsig (32).

### Evidence of stepwise slippage events

#### Sequence composition and slippage events

We developed in-house scripts to count the indel mutations at different lengths, and to extract the motif size & motif repeat units from the Variant Call Format (VCF) files obtained from Hartwig medicine foundation. We calculated the mean frequency of each motif for these indel mutations at different lengths.

Shared unique MS mutations are defined as indels occurring at the same chromosome and position in paired biopsies collected from the individual patient, focusing on MS loci detected with a single indel mutation. The percentage of shared MS loci is calculated as the ratio between the number of shared MS indels and the total indel count within the biopsy (Fig. 2b). The slippage size is calculated as the difference in the repeat number of motifs between time point 2 (sample ID marked ‘II’) and time point 1 (sample ID marked ‘I’).

#### Length-specific deletion count and long-deletion intensity

The deletion number at different lengths (from 1 to 5+ bp) are aggregated based on the annotations provided in the 83-channel profile of each sample (Fig. 1g; Supplementary Fig. 7-9). We also compared the accumulation pattern of deletion count by calculating the log_10_ ratio of the aggregated indel counts between ‘old’ and ‘young’ biopsies.

Estimated number of slippage events is calculated using formula ∑*_l_* _ε [1,5’]_(*l* ∗ *N_l_*). In this equation, *N*_*l*_ represents the aggregated deletion mutation count at specific indel length *l* (where *l* εg [1,5]) from the indel profile with observed mutation count. The length of the deletion (*l*) is used as an approximate measurement of the total slippage event that occurred at the mutated MS loci, as most of the deletions are mononucleotide repeats (Fig. 2e). Similarly, long-deletion intensity is calculated using formula ∑*_l_* _ε [1,5’]_(*l* ∗ *f_l_*). In this equation, *f*_!_ represents the aggregated deletion frequencies at specific indel length *l* (where *l* ε [1,5]) from the normalised indel profile.

### Survival analysis

In the overall survival analysis, the patients’ survival status is determined using recorded clinical information (the ‘Death’ column; *n*=100) from the Hartwig cohort. The survival duration is calculated as the difference between the biopsy date and the ‘DeathDate’ (if the patient is deceased). For patients who are alive, we used the ‘treatmentEndDate’ (if available) or the last updated date in the clinical file (‘2017-07-15’).

We also conducted a disease progression-free survival analysis on 39 cancers in the Hartwig cohort with documented treatment information. For these cases, the event is defined as whether the disease progressed after the initial treatment. Samples labelled ‘PD’ were considered progressors, while those labelled ‘SD’ or ‘PR’ were considered non-progressors. The duration is calculated as the time difference between the ‘responseDate’ and the ‘biopsyDate’.

The Kaplan-Meier (KM) curves and the case numbers at risk in this study were derived from the ‘KaplanMeierFitter’ function from the ‘lifelines’ library, based on the ‘event’ and ‘duration’ data. We grouped samples into ‘higher’ and ‘lower’ categories using the median of different metrics as a cutoff, including long-deletion intensity, TMB (total number of SBSs and indels), and the frequency of 1 bp A/T deletions at HP regions. The statistical significance of the differences between the ‘higher’ and ‘lower’ subgroups in each KM curve was assessed using the ‘pairwise_logrank_test’ function from the ‘lifelines.statistics’ library in Python.

### Feature selection in sWGS samples

We employed a previously published feature selection method (30) to identify significantly informative features for MMRd classification. Firstly, we obtained the true feature importance of the 83 indel signature channels using the original datasets with correct labels of MMR status. We then generated *n=*50 randomly permuted datasets with shuffled labels and computed the feature importance distribution for each signature feature (also termed as ‘null importance’ distribution). The feature score is the log ratio of the true feature importance to the 95% quantile value of the null-importance distribution. We repeated the above process 20 times on the different training sample sets using random five-fold splits from the original dataset. This feature selection process was independently conducted for FFPE and FF cohorts.

### Noise correction in low-purity sWGS samples

To mitigate the noise in MMRd indel profiles with low tumour content, we employed our previously developed tool, FFPEsig (32). We derived the background noise profile by computing the average indel profiles from MMRp samples in the EPICC cohort. Subsequently, the observed indel profile of low-purity MMRd samples along with the derived noise profile served as input for FFPEsig to predict signal profile and the weight of signal. To address potential noise contamination in longitudinal IBD MMRd samples (Fig. 6c), we applied a similar approach where high-confidence MMRp samples from the same FFPE cohort served as the noise profile. This noise correction step, now available as an optional feature within MILO, allows users to apply either our default or a customised noise pattern to their own samples.

### Development of MILO

#### Simulation of sWGS indel profiles

Given the limited number of real sWGS samples, especially MMRd cases with low tumour purity, we simulated an additional 10,000 MMRd and 10,000 MMRp samples with varying purity levels to train automatic classifiers for MMRd detection. Out of 10,000 MMRd cases, 5,000 samples had relatively low purity (<0.2).

To generate synthetic MMRd samples, we mixed MMRd (the ‘signal’) and MMRp (the ‘noise’) profiles together. The signal profile was based on the averaged high-purity (>0.9) MMRd profiles from the EPICC cohort, while the noise profile was randomly selected from MMRp samples in the FFPE or FF datasets. The noise indel count remained nearly constant, with ∼42,100 and ∼127,839 noise mutations for FF and FFPE samples, respectively, at 1X coverage (Fig. 5d-f). The signal mutation count was predicted using the linear regression model as trained in Fig. 5e, with tumour purity as the input variable. A simulated indel profile with purity *i* (randomly chosen from a uniform distribution) was then created by multiplying the signal profile by the signal mutation count and adding it to the product of the noise profile and the noise mutation count. We added Poisson noise to the noise & signal levels as well as to each 83 indel profile channels. For simulating MMRp samples, no MMRd signal was added, and the rest of the procedure remained the same.

#### Developing and testing MILO

Firstly, we generated a set of simulated samples only using MMRp samples randomly selected from two out of three FF or FFPE cohorts: T1-CRC and UC-IBD for FFPE, and ‘Gland’ and Mseq for FF. The third cohort (Duodenal for FFPE and EPICC for FF) was reserved for independent testing to assess whether classifiers trained on the simulated samples could accurately predict MSI status in an unrelated cohort.

To select the most effective approach, we trained three classifiers: RandomForestClassifier (RF) (51), GaussianNB (NB) (52,53), and Support Vector Machine (SVM) (54)—from the ‘sklearn’ package on the simulated samples. Their performance was evaluated using 5-fold cross-validation. False positive and true positive rates were generated using the ‘roc_curve’ function in ‘sklearn’, and the Area Under the Curve (AUC) was calculated for each Receiver Operating Characteristic (ROC) curve. This process was repeated 50 times with different random seeds. Our results showed that the random forest classifier performed best. Therefore, we used the trained random forest model to predict MMR status in the third independent FF or FFPE cohort, with the predicted results and true labels used to derive the confusion matrix (Supplementary Fig. 25).

Next, we re-simulated a new set of 20,000 samples using MMRp samples from all three FFPE/FF cohorts. We trained two Random Forest classifiers—one for FF samples and one for FFPE samples—on these synthetic and real low-pass samples. These classifiers are now implemented in MILO, and they provide the probability of each sample being an MSI, termed as Prob(MSI). In MILO, the categorical classifications are based on Prob(MSI): ‘non-MSI’ (<=0.5) ‘MSI’ (>=0.75; high-confidence) and ‘Maybe’ (>0.5 & < 0.75; low-confidence). Additionally, MILO can optionally estimate and report MSI intensity of predicted MMRd samples using their noise-corrected indel profiles.

Finally, we compared MILO performance on low-purity samples to that of a widely used MSI detection tool, MSIsensor2 (33), following their recommended protocol with ‘-c 1’ in the command line. Samples with a minimum ‘somatic_site’ percentage of 20% were considered MSI-positive.

## Data Availability

Processed data needed to recreate all results will be available in the Supplementary. This study used VCF files generated from deep-sequenced MMRd and/or MMRp samples from publicly available data cohorts, including EPICC (19,20), PCAWG (13,21), GEL (22,23), Hartwig (24) and *MSH6* knockout experiment (12). We followed the data accession protocols described in the original publications to retrieve the necessary data. Sequence data for EPICC samples have been deposited at the European Genome-phenome Archive (EGA), which is hosted by the European Bioinformatics Institute and the Centre for Genomic Regulation, under accession number EGAS00001005230. Additionally, we incorporated several publicly available sWGS cohorts into our analysis. These cohorts can be downloaded through the data descriptions in their respective original publications: UC-IBD (27), T1-CRC (28), EPICC_lpWGS (19,20), Gland_lpWGS (28) and Mseq_ds_lpWGS (29). Note that sWGS BAMs for the Mseq_ds_lpWGS cohort were generated using a down-sampling approach from the original deep-sequenced BAM files with an EGA accession number of EGAS00001003066. BAM files from duodenal samples are in the process of being submitted to the EGA database (accession number to be provided soon). Raw sequencing data of longitudinal IBD samples is not available due to the retrospective and anonymous nature of the study.

## Code Availability

MILO is available from https://github.com/QingliGuo/MILO/blob/main/README.md. The analysis code for reproducing results will be made available on Github.

## Contribution

Q.G - conceived the study, conceptualisation, data curation, data analysis, visualisation & presentation, bioinformatics pipeline construction, interpretation, writing, review & editing. J.H - GEL & EPICC data analysis, Hartwig data curation, interpretation, review & editing. E.L - interpretation, presentation, editing. S.N - duodenal cancer data curation, CNV calling, data interpretation & editing. I.AB - IBD sample collection and NGS data generation. H.G - phylogenetic tree construction & interpretation, editing. V.B - duodenal cancer data generation & review. C.S.H - duodenal cancer data generation & review. L.Z - Slippage event analysis interpretation & editing. H.M - duodenal cancer sample generation, review & editing. A.S - EPICC data curation, interpretation & editing. A.B - sample curation, interpretation, data presentation, review & editing. V.M - conceived the study, conceptualisation, funding acquisition, interpretation, supervision, review & editing. T.A.G - conceived the study, conceptualisation, data generation & accession, interpretation, supervision, writing, review & editing. All authors read and approved the final manuscript.

## Acknowledgements

QG received a postdoctoral fellowship (Schottlander Cancer Breakthrough Award) from the Schottlander Foundation via the Institute of Cancer Research. QG was in part supported by the University of Helsinki and the Research Council of Finland (Grant 345829 to VM). TG received funding from Cancer Research UK (DRCNPG-May21_100001; A19771), the Wellcome Trust (202778/Z/16/Z) also with AS, the Barts Charity (472–2300) and the NIH (U54 CA217376) also with AS, for various aspects of this work. H.M acknowledges the support from Barts Charity, Barts Health NHS Trust and Cancer Research UK Core grant for generating the duodenal cohort (UKDCSG). EL is supported by a start-up grant from the Chalmers Area of Advance Health Engineering. We thank Dr. Ignacio Vázquez-García, Dr. Virinder Singh Reen, Dr. Frederick J H Whiting and Dr. Calum Gabbutt for their constructive comments and discussion on the manuscript. We acknowledge the QMUL core facilities for their sequencing and histology support.

## Supplementary Figures

**Supplementary Fig. 1.**
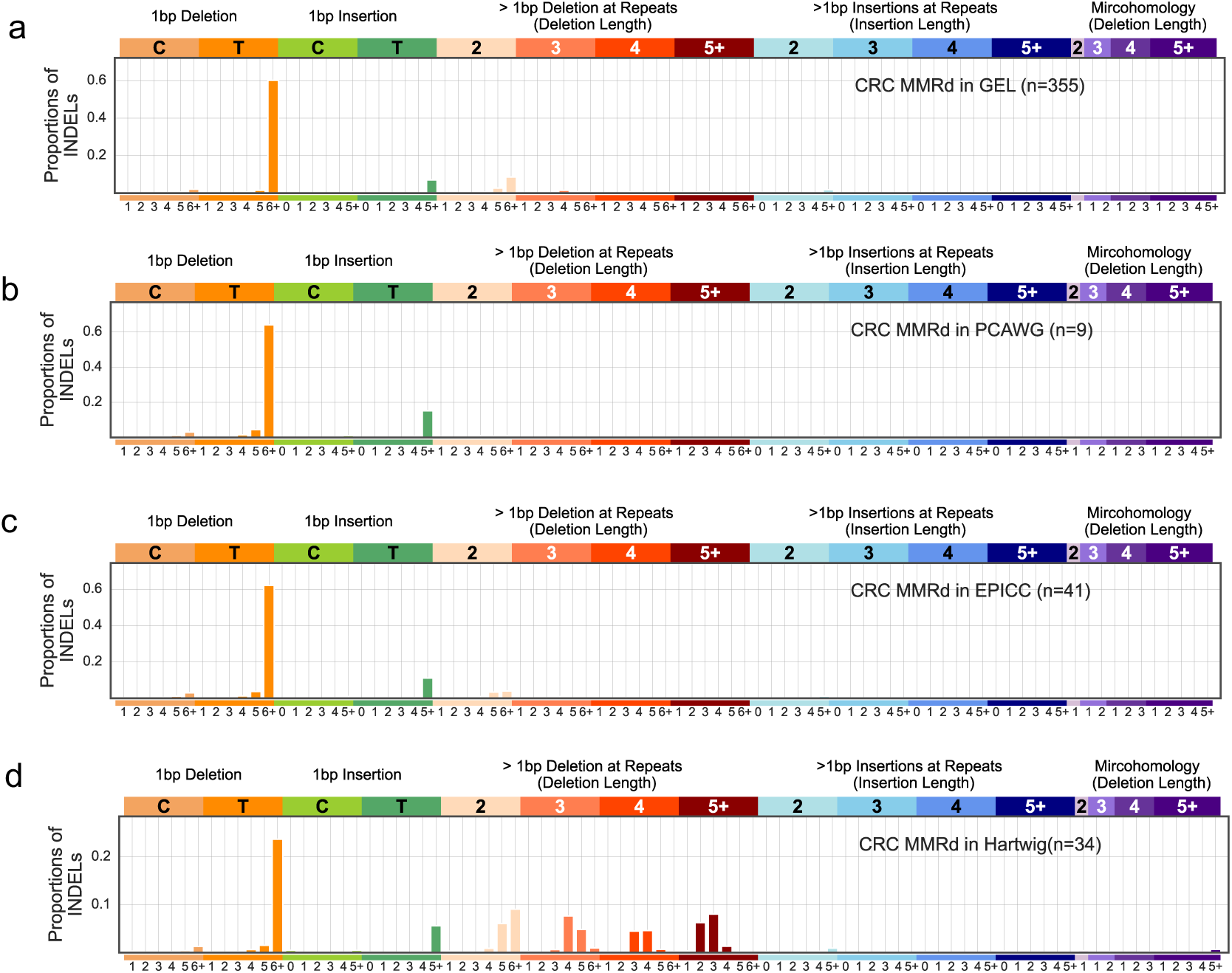
The mean indel profiles of CRC MMRd samples in four deep-sequenced data cohorts. We normalised the average indel profiles of CRC MMRd cancers from (**a**) GEL, (**b**) PCAWG, (**c**) EPICC and (**d**) Hartwig cohort.

**Supplementary Fig. 2.**
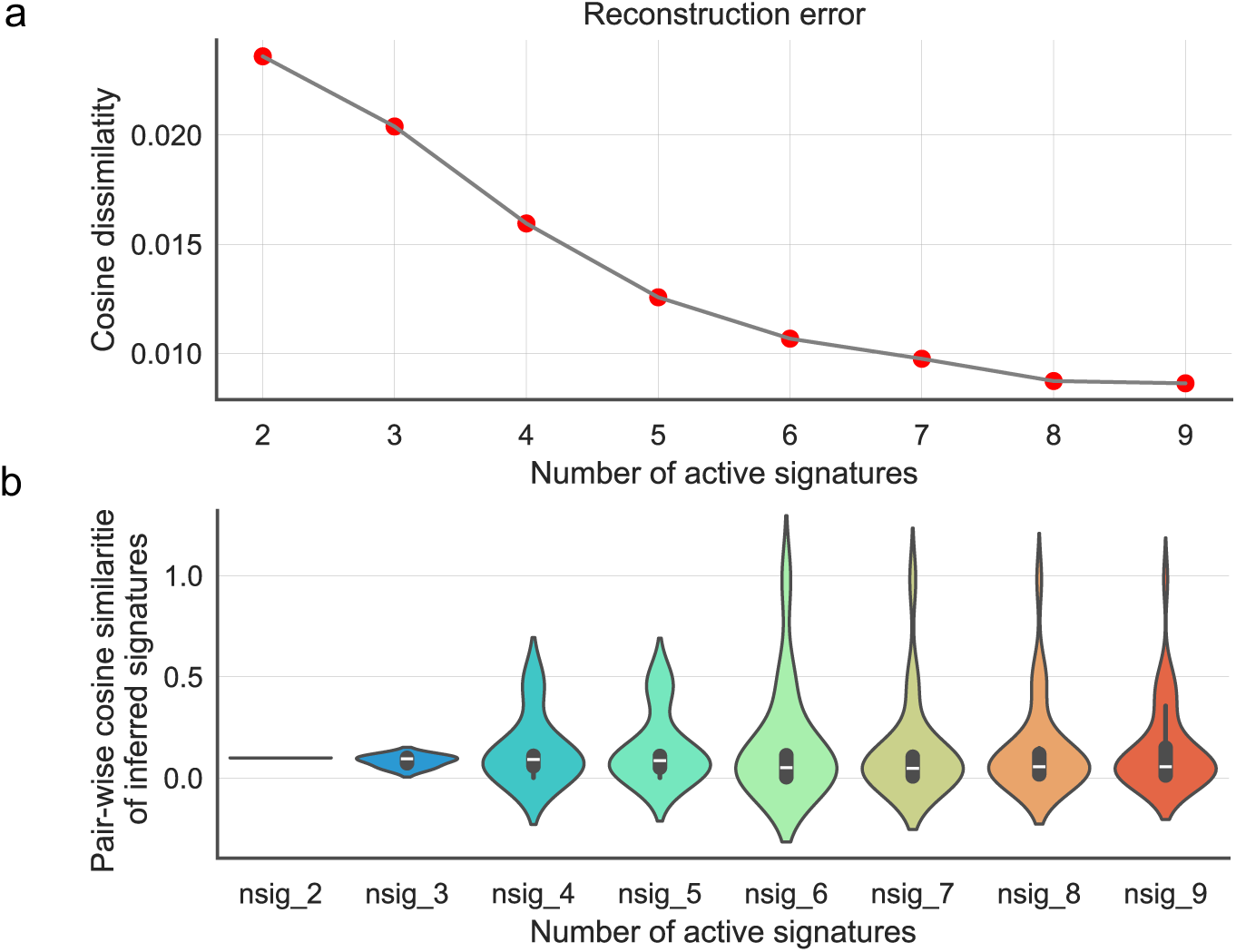
Model selection for the number of active indel signatures in deep-sequenced MMRd cancers. (**a**) The cosine dissimilarity between the original and reconstructed indel profiles, and (**b**) the pairwise cosine similarity between inferred signatures. The width of the violin plot reflects the frequency of data points in each region, and the overlaid box plot in the centre indicates the interquartile range, with the white line representing the median.

**Supplementary Fig. 3.**
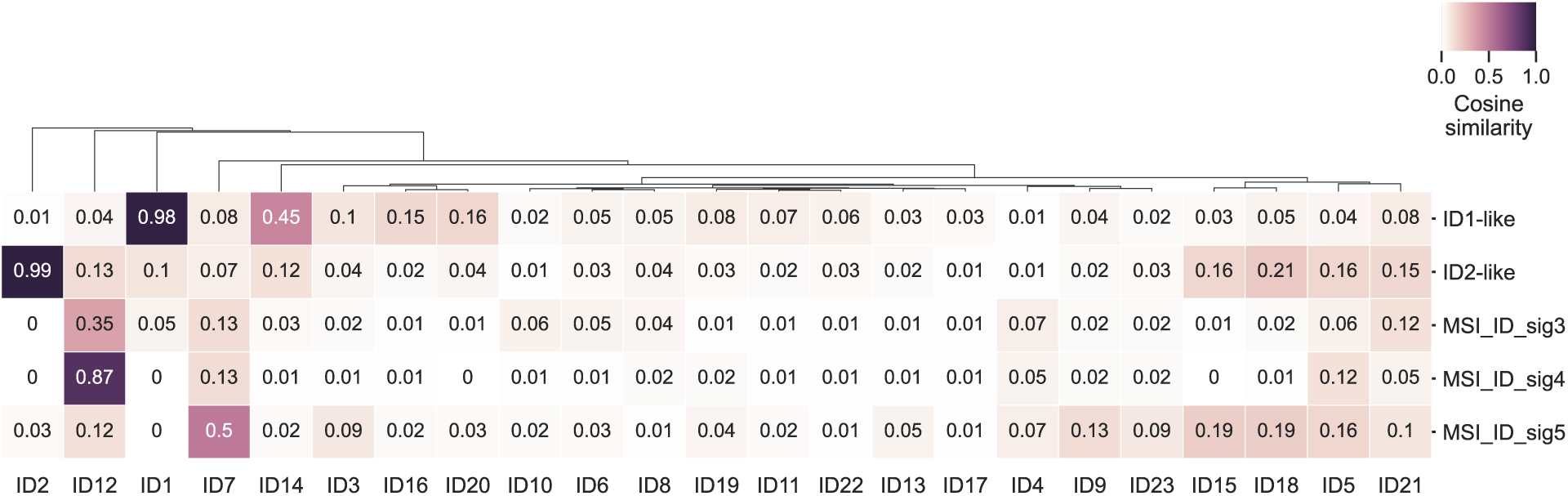
Comparison of inferred MSI indel signatures to known indel signatures. Known indel signatures (COSMIC v3.4) are displayed in columns, while the five newly inferred MSI indel signatures are shown in rows. Cosine similarities for each comparison are annotated in the heatmap.

**Supplementary Fig. 4.**
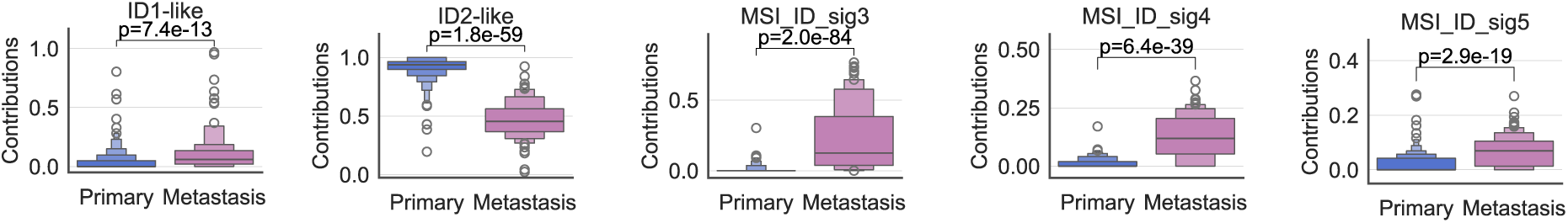
Relative contributions of inferred MSI indel signatures in primary and metastasis MMRd cancers. Data are presented using a Letter-Value plot and the black line corresponds to the median of the dataset and every further step splits the remaining data into two halves. Statistical significance was assessed using a two-sided Mann-Whitney *U* test.

**Supplementary Fig. 5.**
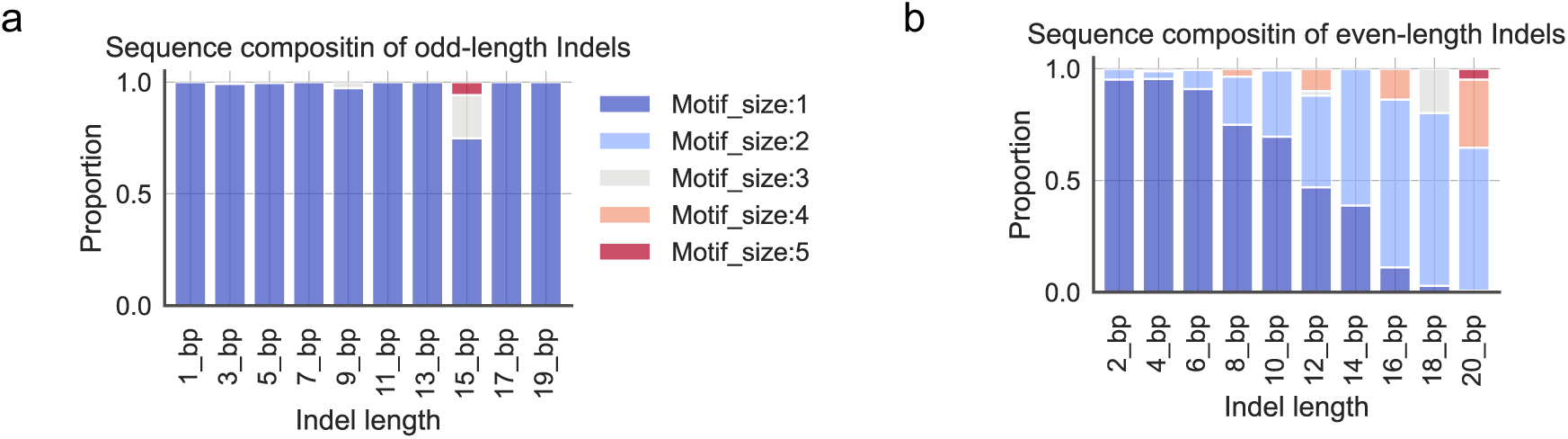
Sequence composition of indel mutations by length in paired MMRd samples from the Hartwig cohort. (**a**) Indels with odd-length (**b**) and indels with even-length. For indels of specific lengths, we calculated the percentages of indels with different motif sizes (ranging from 1 to 5). The percentage data from the ten biopsies were normalised and presented as bar heights.

**Supplementary Fig. 6.**
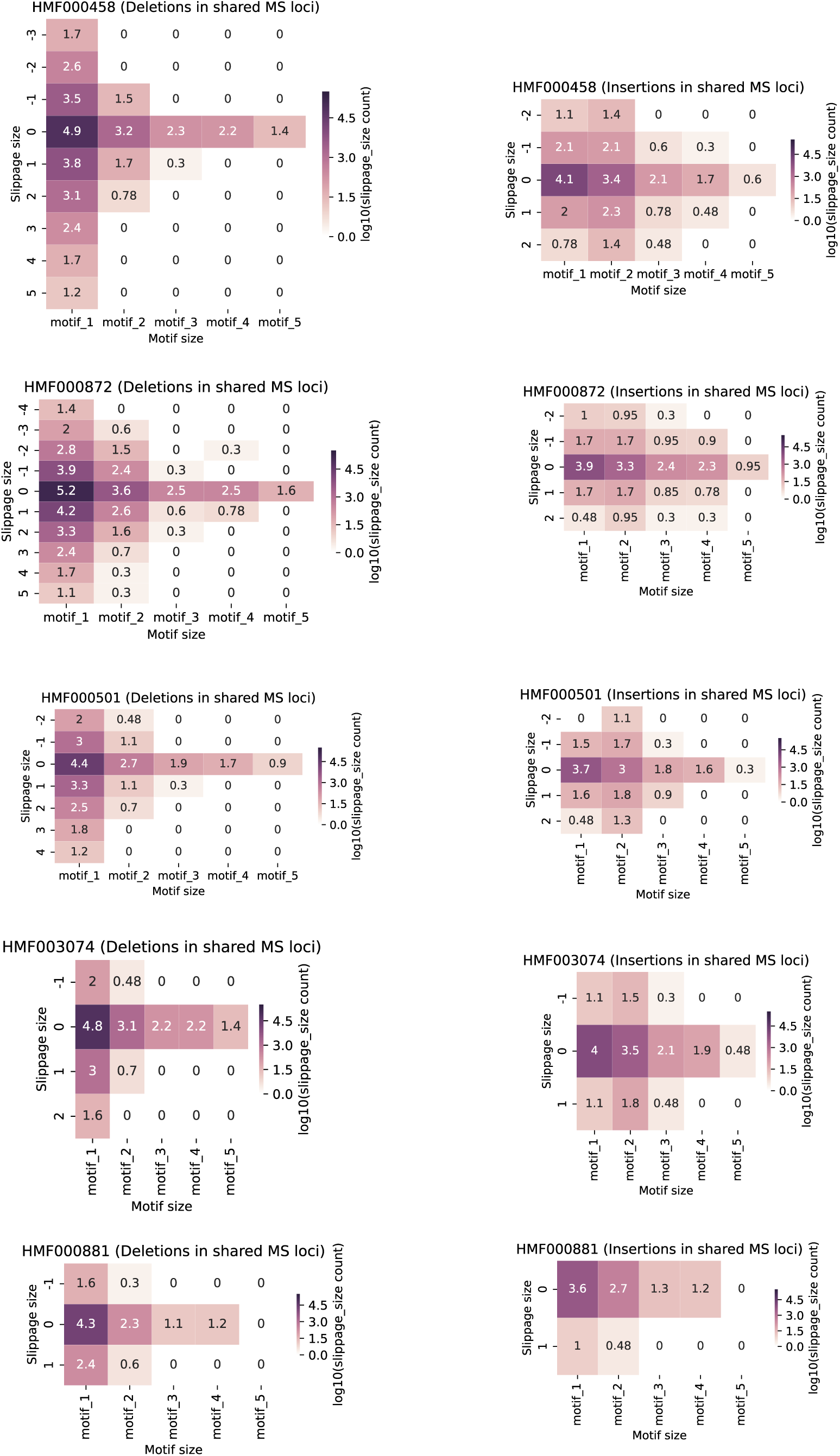
Heatmaps showing the number of slippage events in ten longitudinal MMRd samples from Hartwig cohort. The heatmaps display the log_10_(number of slippage events) according to slippage size (Y-axis) and motif size (X-axis).

**Supplementary Fig. 7.**
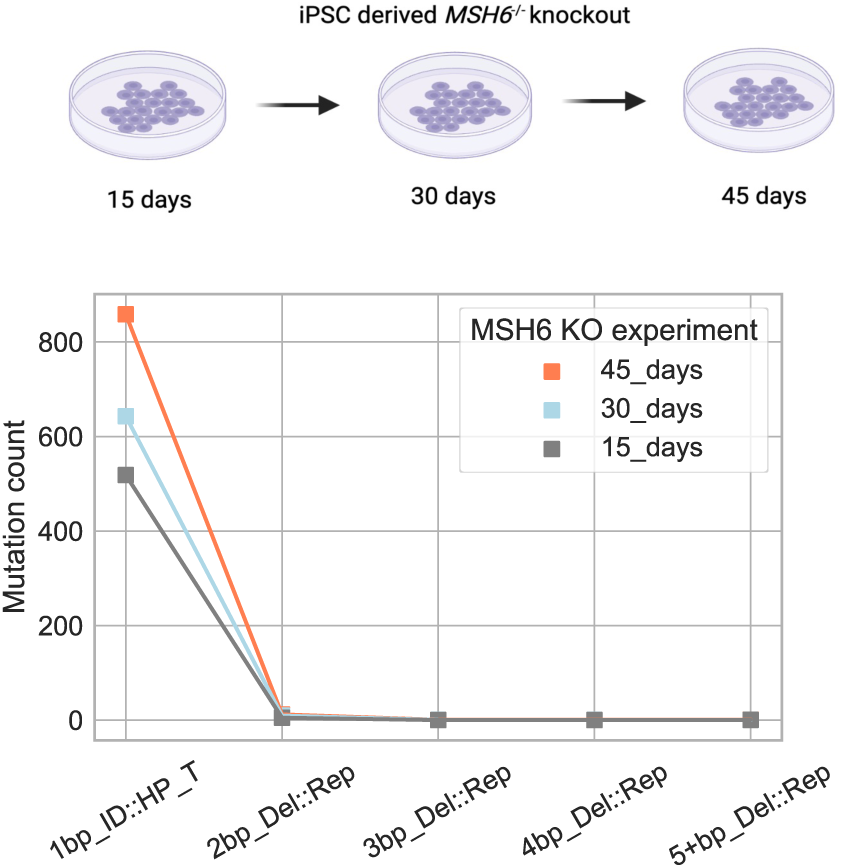
Deletion counts by length in longitudinal samples from *MSH6 knockout (*KO) cell lines. A total of 8 samples were collected and deeply sequenced WGS: two samples each at 15 and 30 days, and four samples at 45 days. We normalized the deletion counts by length for each time point.

**Supplementary Fig. 8.**
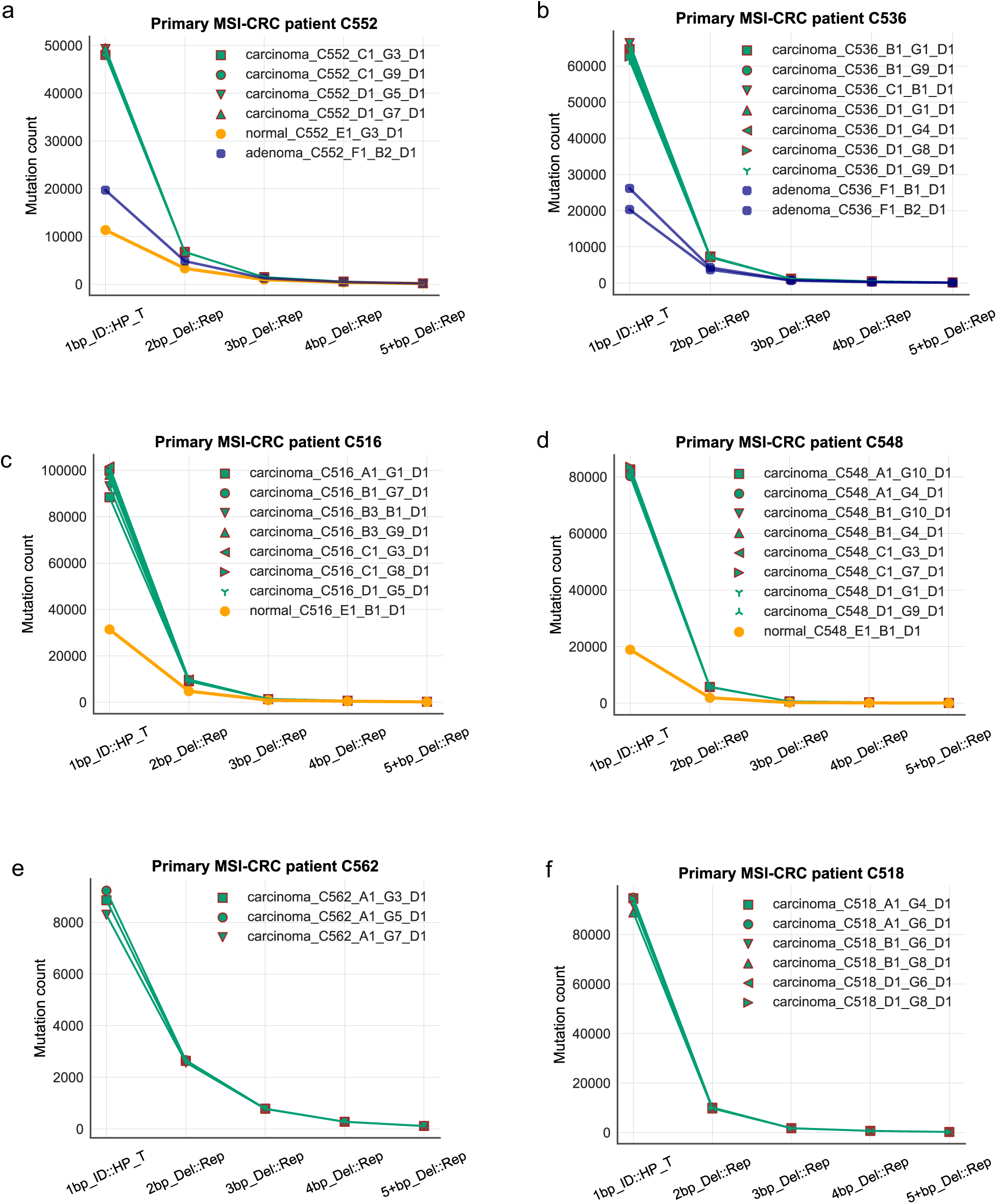
Deletion counts by length in multi-region samples from six primary MMRd CRC patients, C552 (**a**), C536 (**b**), C516 (**c**), C548 (**d**), C562 (**e**) and C518 (**f**). The multi-region biopsies (deep-sequenced) are available for each patient which were categorised into three main histological subgroups with different coloured dots (carcinoma: green; adenoma: blue; normal-looking: golden). All biopsies for each patient are annotated with different shapes and biopsy IDs, both of which are indicated in the legend of each subfigure.

**Supplementary Fig. 9.**
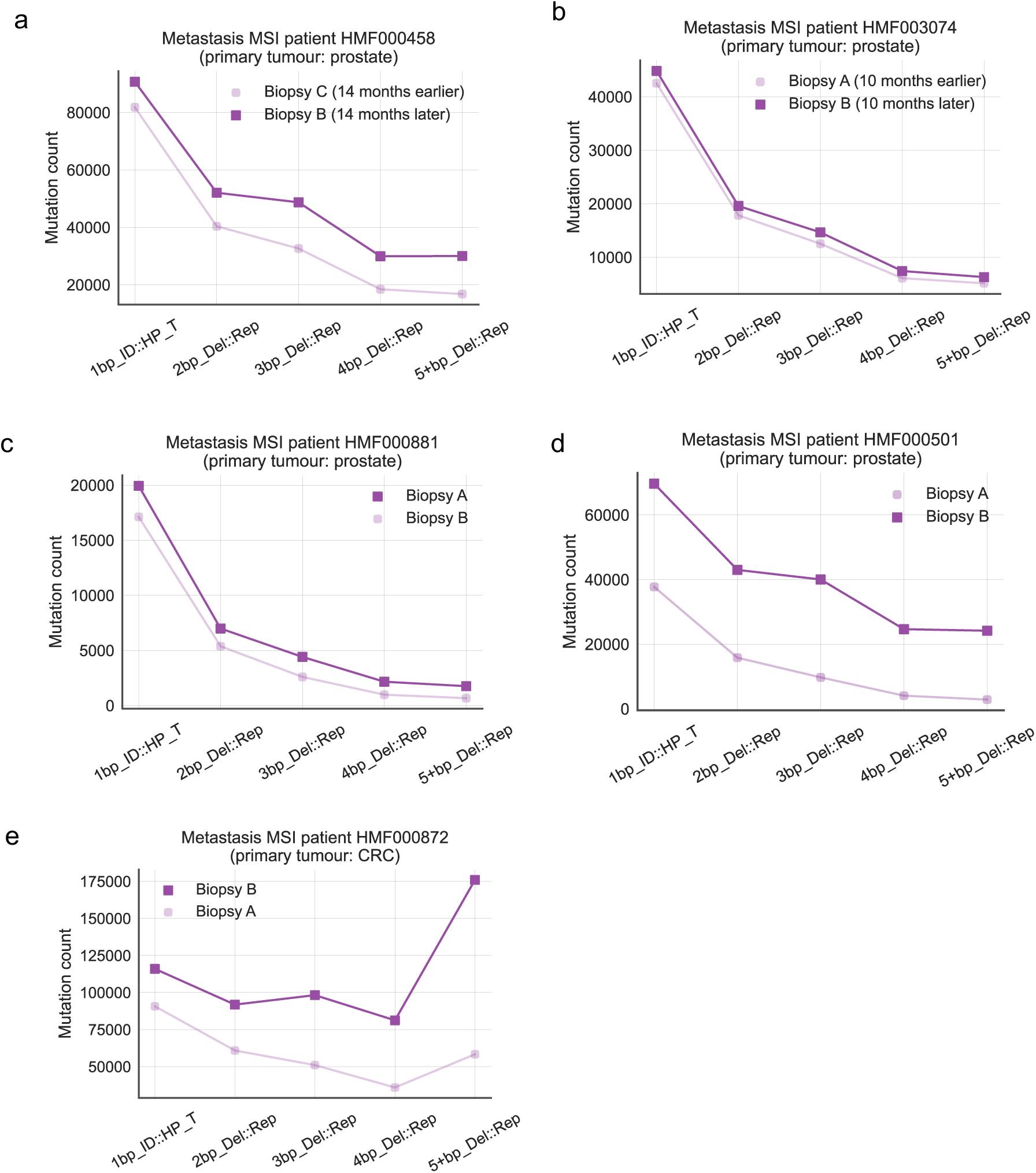
Deletion counts by length in longitudinal samples from five metastasis MSI patients (Hartwig). Two longitudinal samples are available for each MSI patient, distinguished by light and dark purple dots. (a-b) Patient HMF000458 (a) and HMP003074 (b) have biopsy intervals of 14 months and 10 months, respectively. (c-e) Patient HMF000881 (c), HMF000501 (d), and HMF000872 (e) have no documented biopsy intervals.

**Supplementary Fig. 10.**
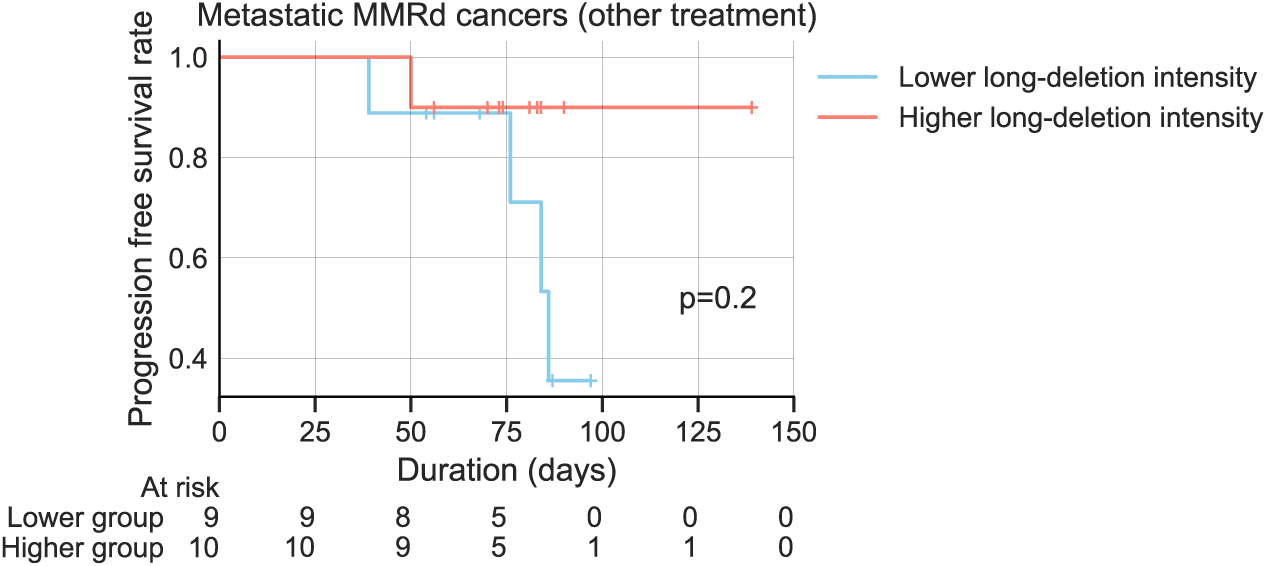
Kaplan-Meier progression-free survival (PFS) analysis for 19 metastatic MMRd cancers. All samples were from Hartwig cohort and received non-chemotherapy treatment. They were stratified based on long-deletion intensity (Methods). P-values were obtained from pairwise log-rank tests.

**Supplementary Fig. 11.**
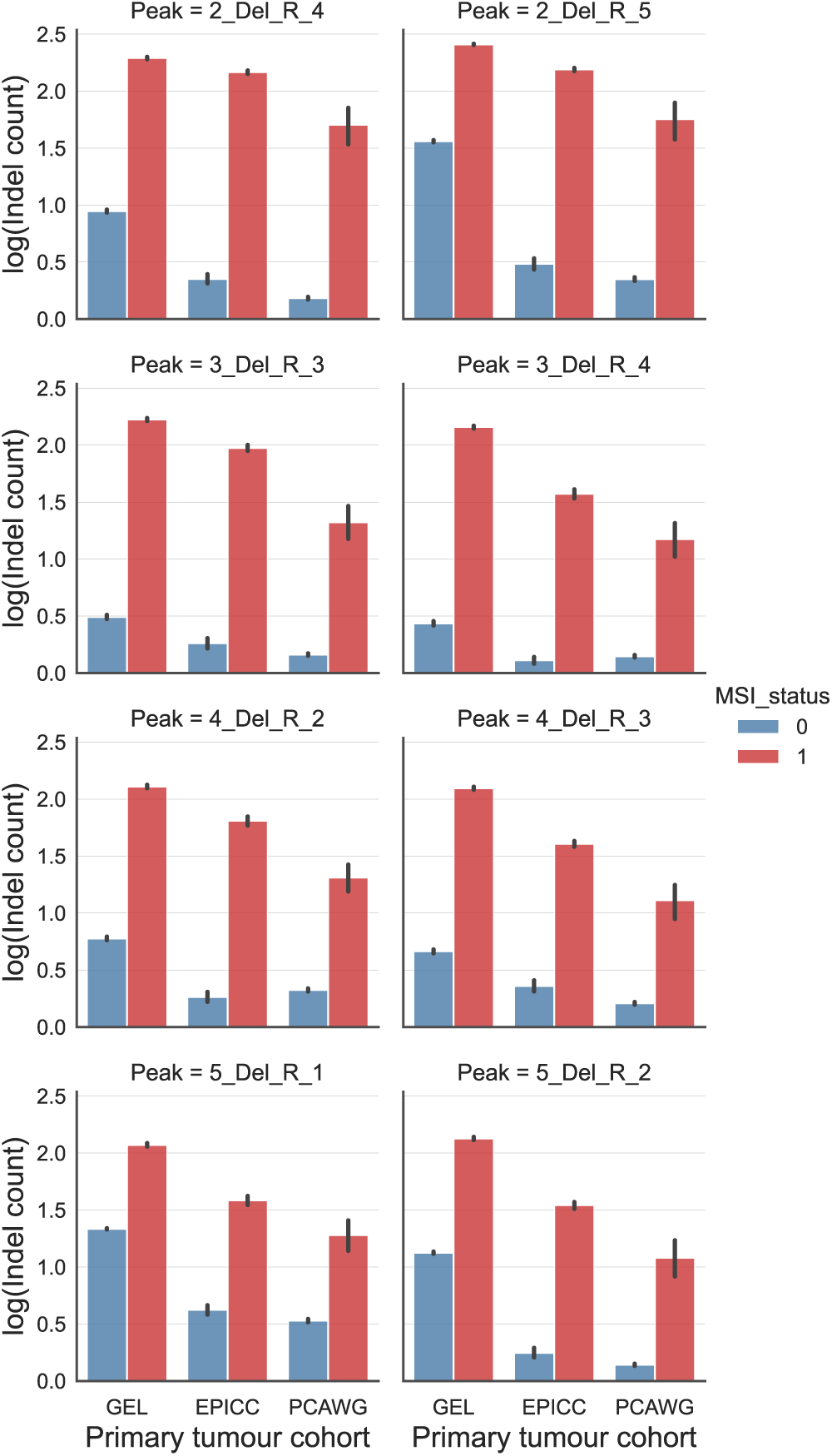
Comparison of long-deletion signature peaks between MMRd and MMRp samples in deep-sequenced primary cancers. Signature peaks refer to long-deletion signatures channels with a mutation frequency greater than 0.05. The *Y*-axis shows the log-transformed counts, and the *X*-axis indicates the data source. The mean values are indicated as the bar heights, with error bars representing the standard deviations.

**Supplementary Fig. 12.**
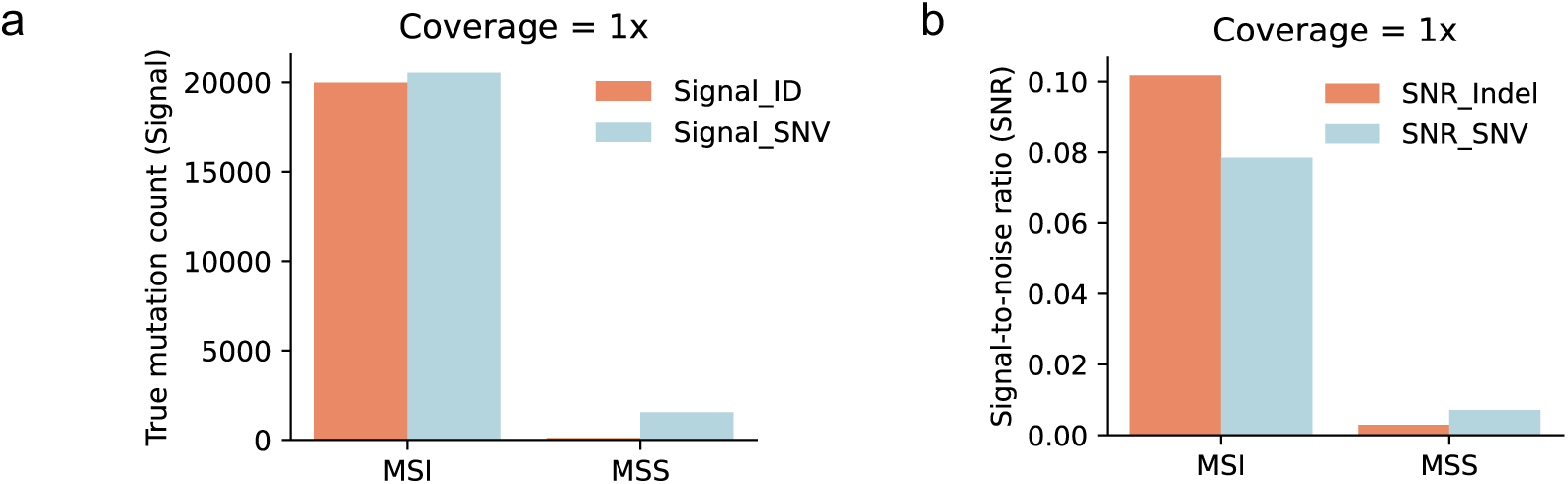
True signal mutations in example MSI and MSS sWGS samples. (**a**) Shows the count of true somatic mutations in MSI and MSS sWGS samples. (**b**) Shows the signal-to-noise ratio in the MSI and MSS sWGS samples. MSS (microsatellite stable) cancers are MMR-proficient (MMRp).

**Supplementary Fig. 13.**
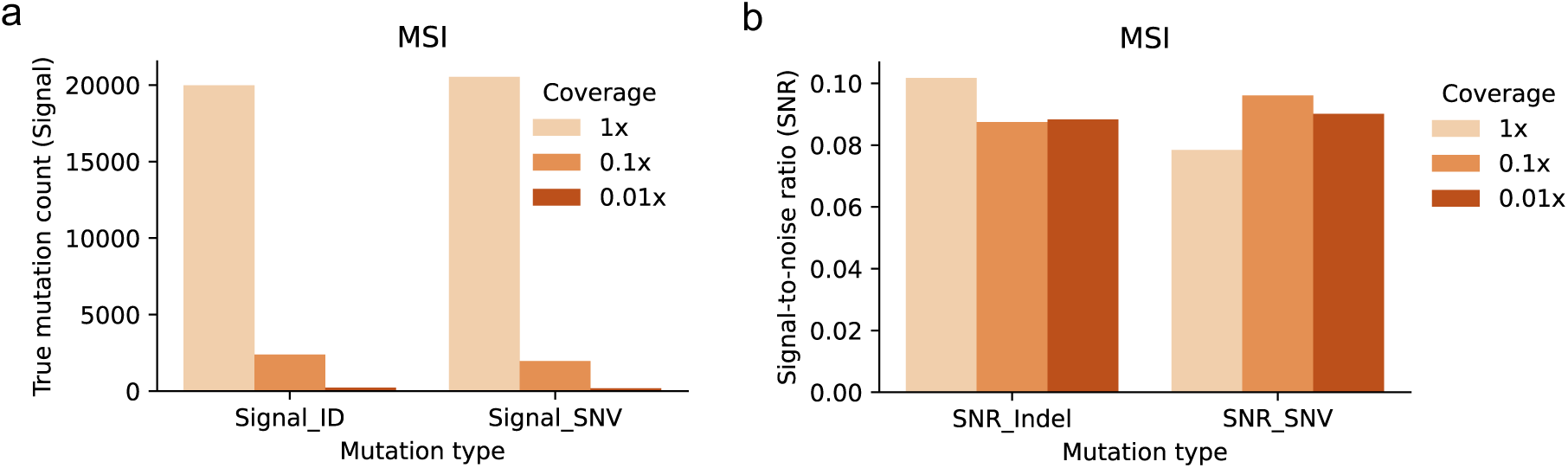
Comparison of true signal mutations in MMRd sWGS samples at different coverages. (**a**) somatic mutation counts and **(b)** signal-to-noise ratios. In addition to the experimentally sequenced sWGS data (coverage of 1X), we generated synthetic sWGS MMRd samples with coverages of 0.1X and 0.01X by down-sampling reads from deep-sequenced data (Methods).

**Supplementary Fig. 14.**
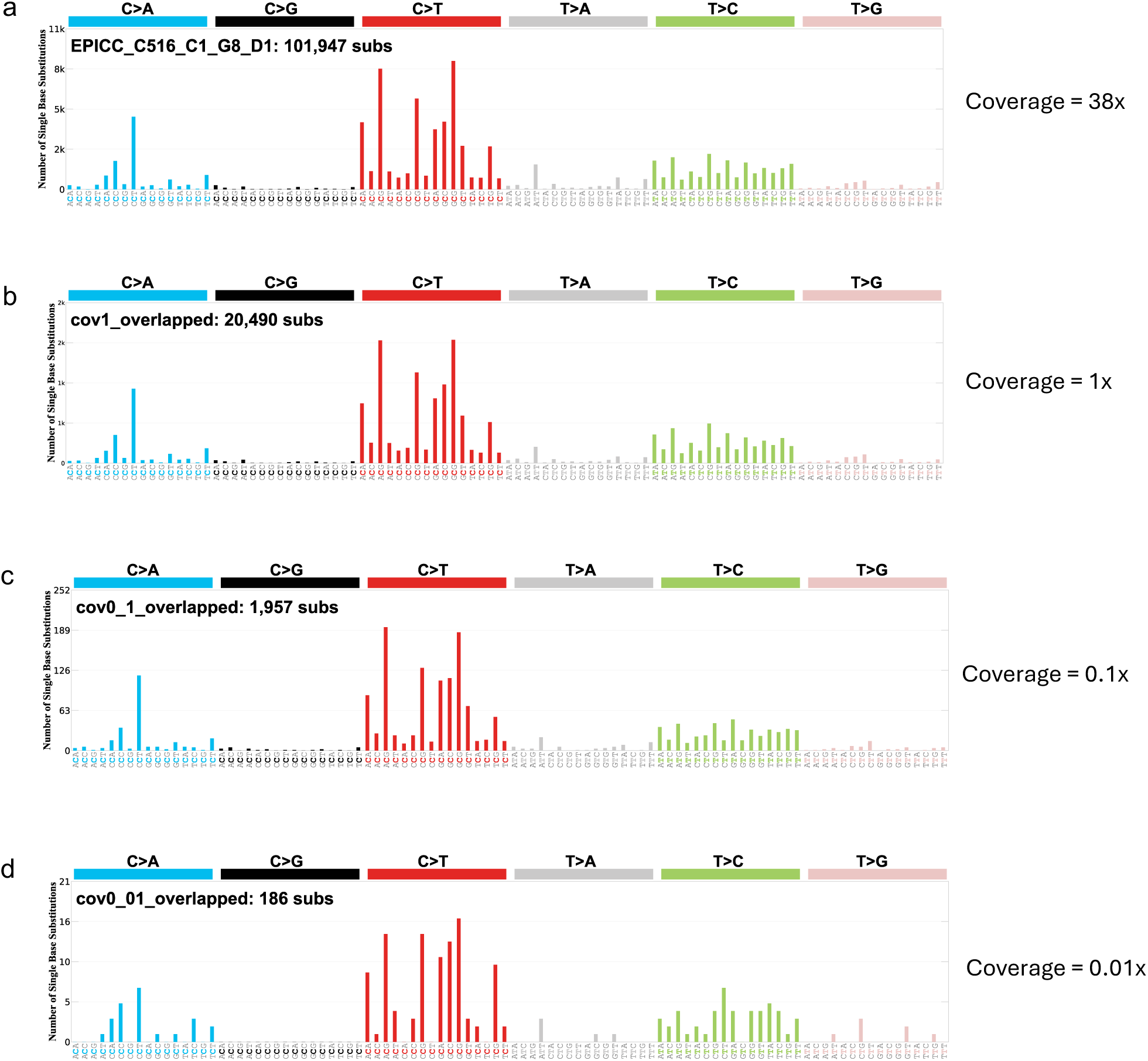
SBS profiles of true signals in an example MMRd cancer at various coverage levels. (**a**) SBS profile derived from somatic mutations identified in a deep-sequenced (coverage of 38X) EPICC MMRd sample with a matched normal. These mutations serve as benchmark mutations. (**b-d**) SBS mutation profiles of true signal SNVs identified in sWGS data at coverages of 1X (**b**), 0.1X (**c**), and 0.01X (**d**), using the benchmark SBS mutations as a reference. Note that matched normal sample was not used for mutation calling in (**b**-**d**) to mimic the scenario of the sWGS samples analysed in Figs. 4-6, where a matched normal is unavailable.

**Supplementary Fig. 15.**
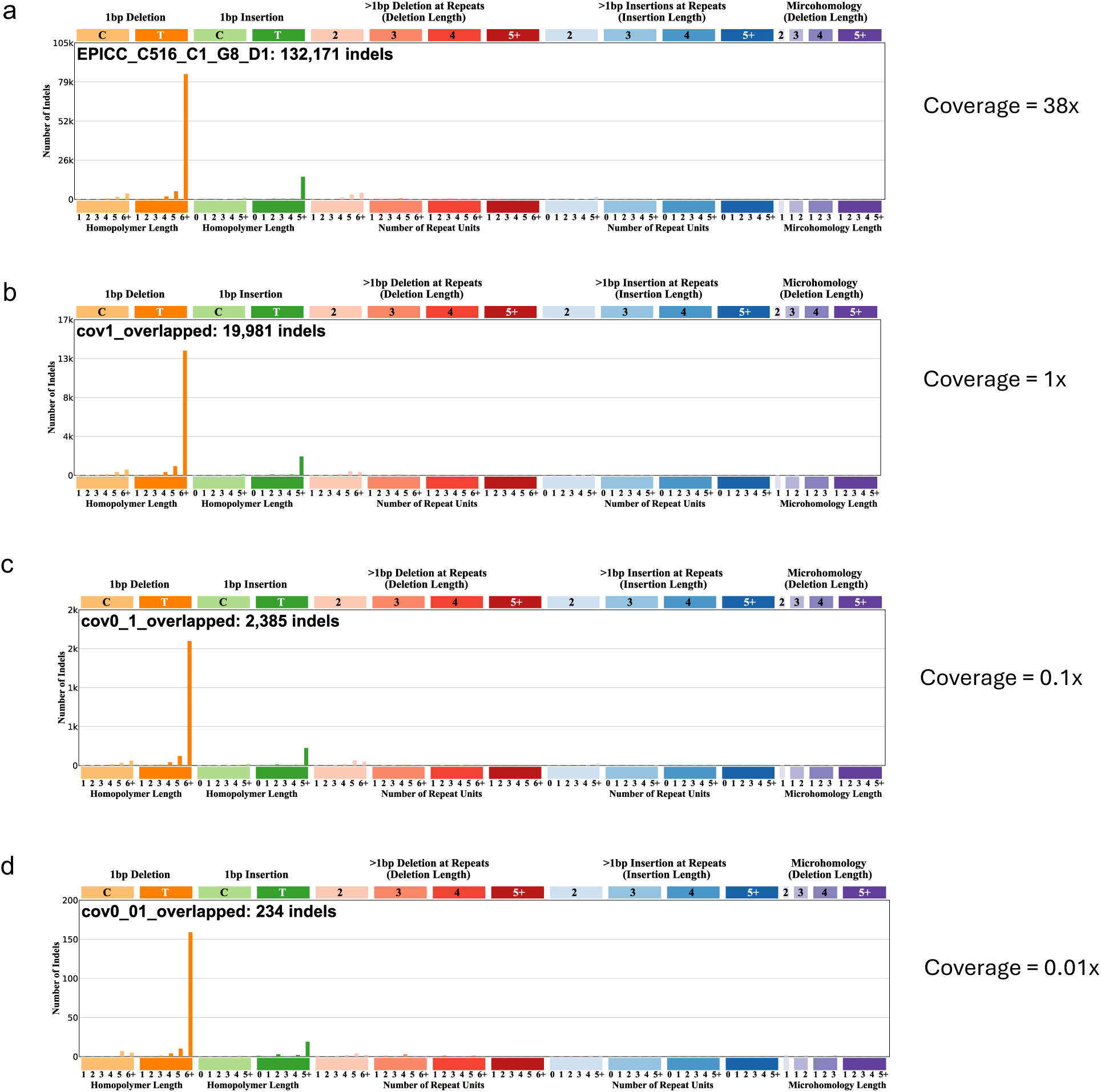
Indel profiles of true signals in an example MMRd cancer at various coverage levels. (**a**) Indel profile derived from somatic mutations identified in a deep-sequenced (coverage of 38X) EPICC MMRd sample with a matched normal. These mutations serve as benchmark mutations. (**b-d**) Indel profiles of true signals identified in sWGS data at coverages of 1X (**b**), 0.1X (**c**), and 0.01X (**d**), using benchmark indel mutations as a reference. Note that matched normal sample was not used for mutation calling in (**b**-**d**) to mimic the scenario of the sWGS samples analysed in Figs. 4-6, where a matched normal is unavailable.

**Supplementary Fig. 16.**
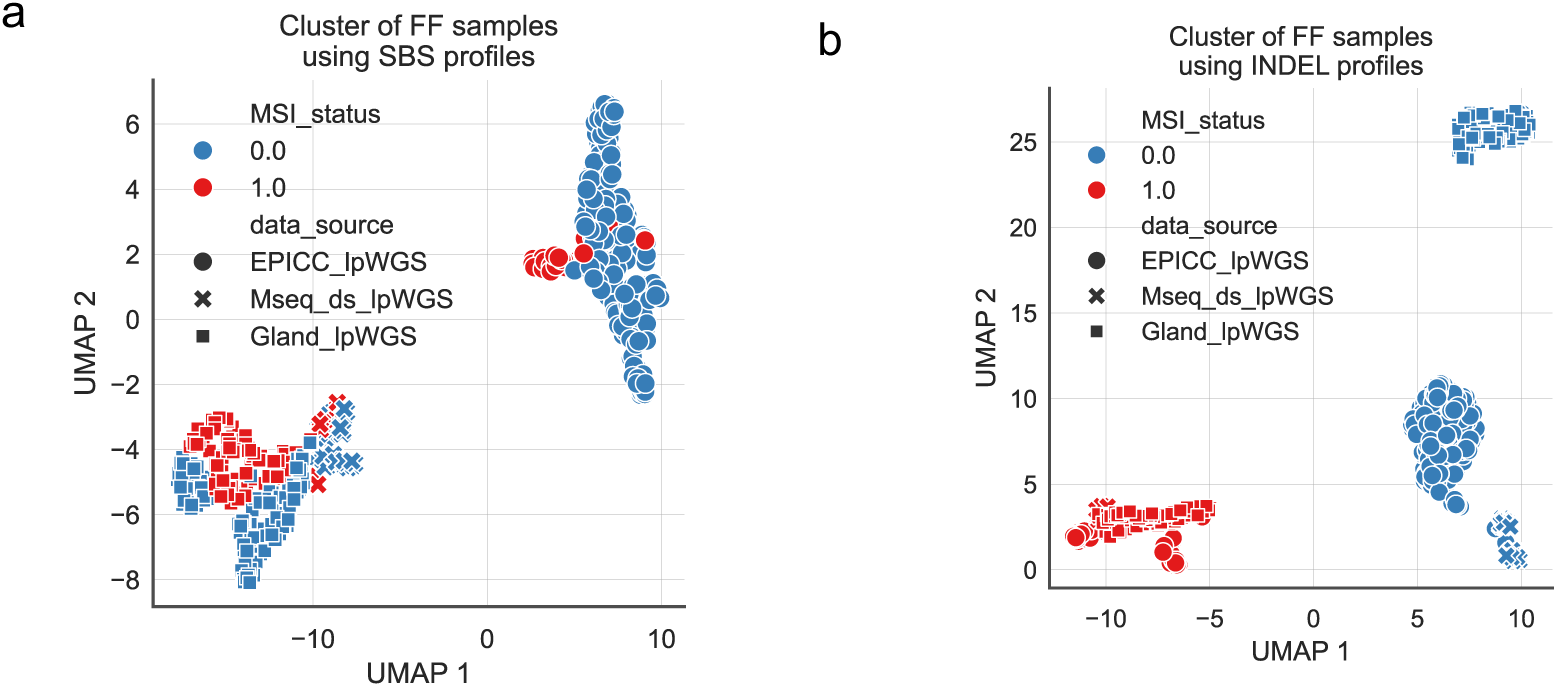
Clustering of FF sWGS samples using SBS (a) and indel (b) mutation profiles. We applied the dimension reduction method, Uniform Manifold Approximation and Projection (UMAP) using either their 96-channel SBS profiles (**a**) or 83-channel indel profiles (**b**). The MSI status of ‘0’ refers to non-MSI/MMR-proficient (MMRp) samples (blue dots), and the MMR status of ‘1’ refers to MSI/MMRd samples (red dots).

**Supplementary Fig. 17.**
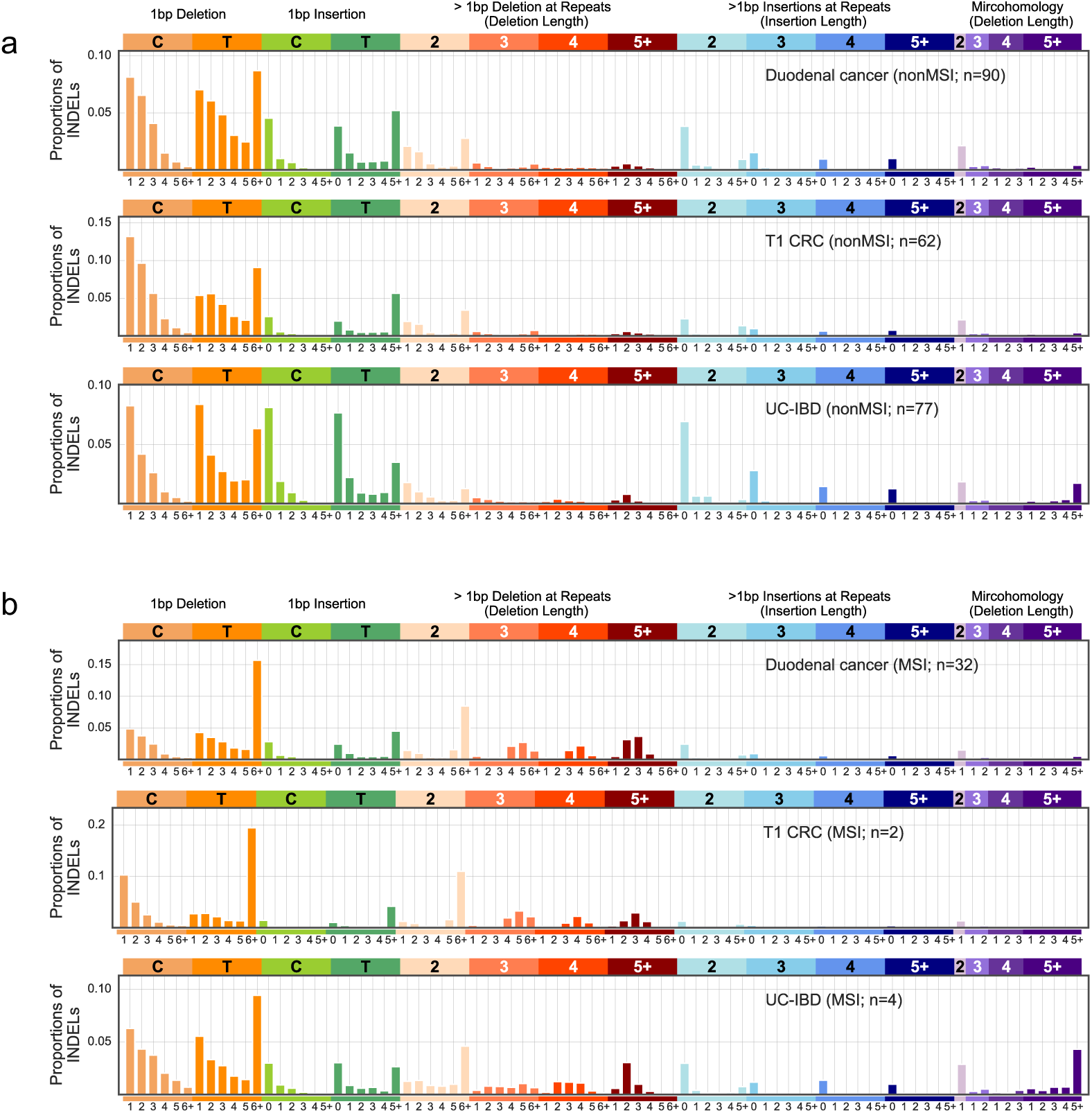
Averaged indel mutation profiles of MMRp (a) and MMRd (b) samples in three FFPE sWGS cohorts.

**Supplementary Fig. 18.**
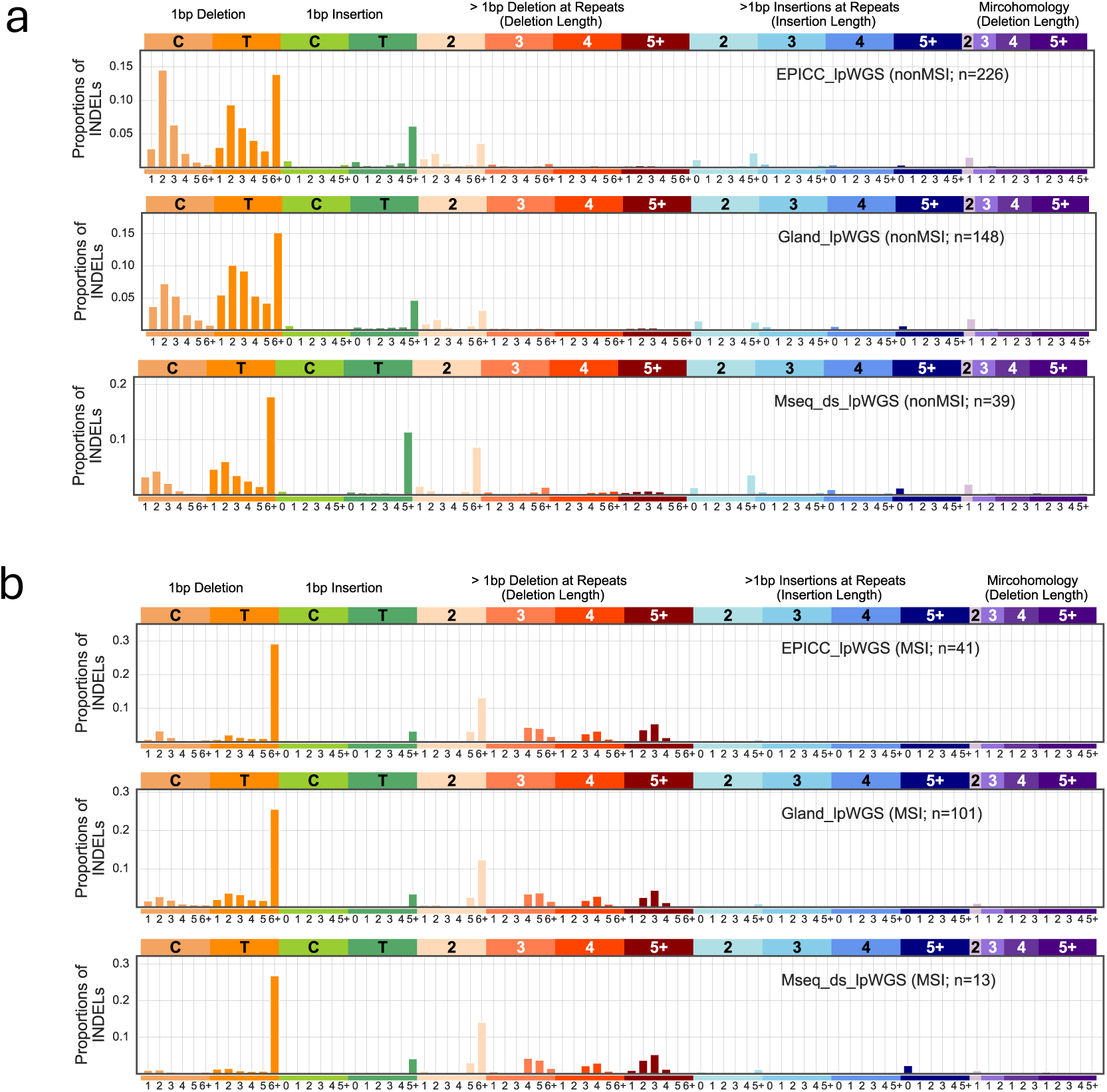
Averaged indel mutation profiles of MMRp (a) and MMRd (b) samples in three FF sWGS cohorts.

**Supplementary Fig. 19.**
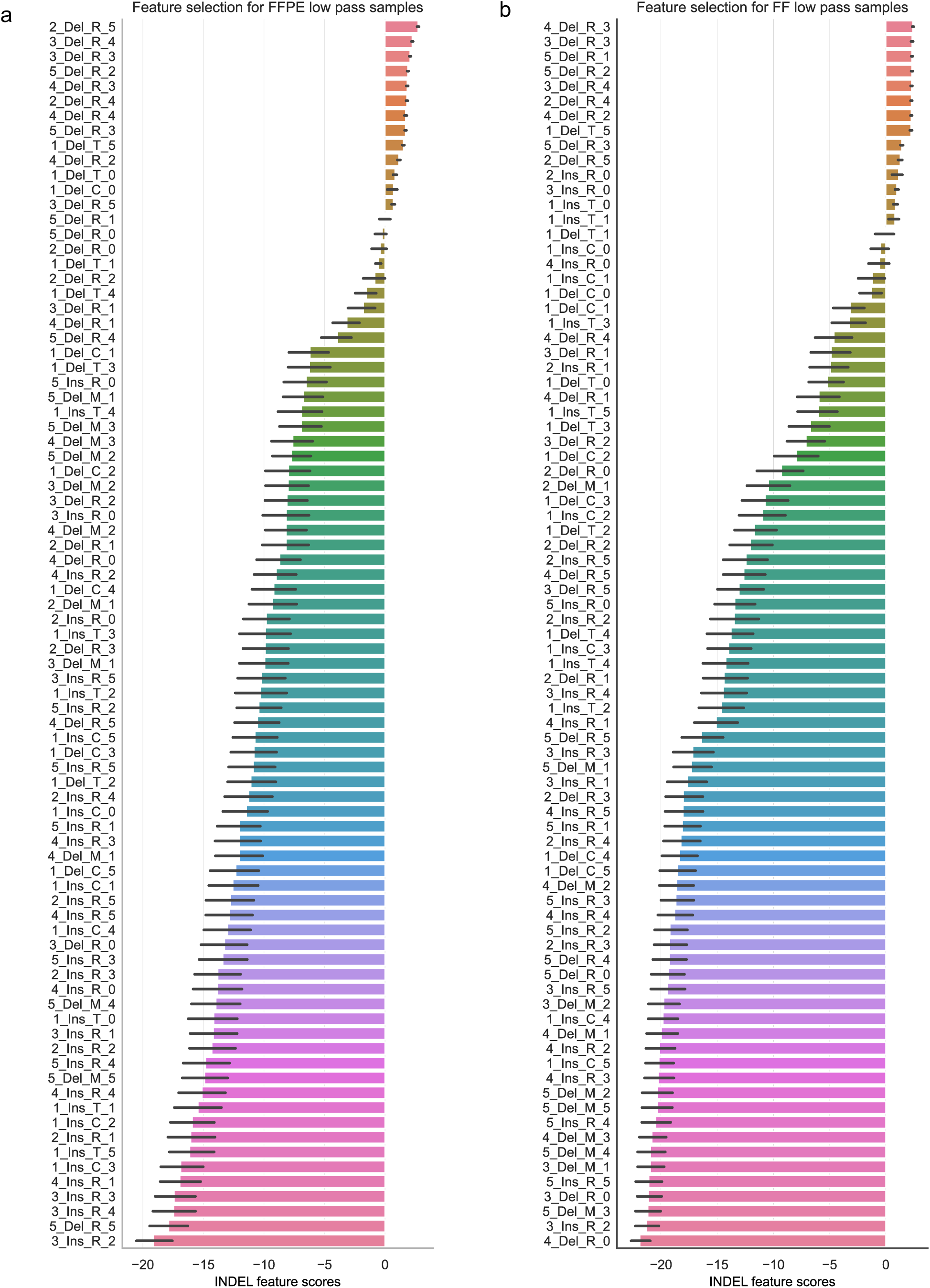
Feature scores of indel signature channels in FFPE (a) and FF (b) sWGS samples. The feature score is calculated as the logarithm of the ratio between the actual feature importance and the 0.95-quantile of the target-shuffled importance score (Methods). We employed a 5-fold sample split, using different randomisations, and summarised the standard deviation of the feature scores from these random splits as the error bar.

**Supplementary Figure 20.**
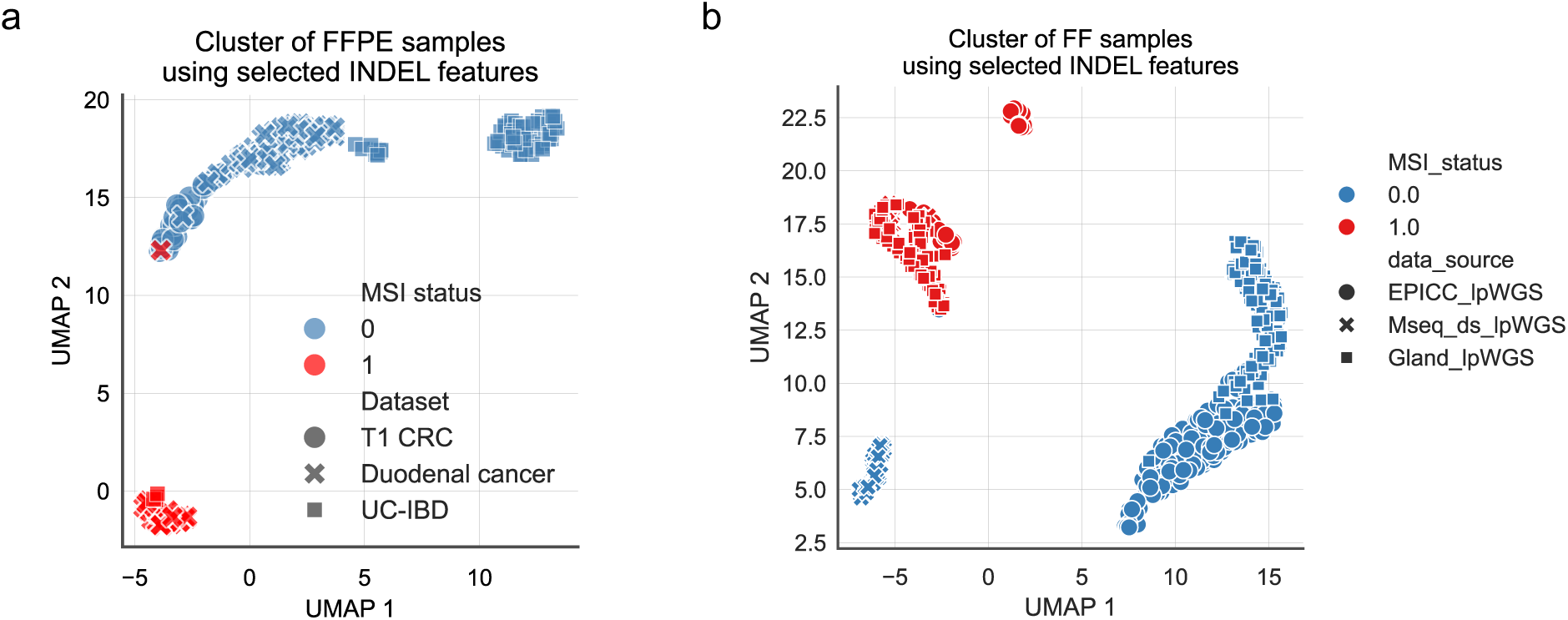
Clustering of FFPE (a) and FF (b) sWGS samples using key indel signature features. We applied the dimension reduction method, Uniform Manifold Approximation and Projection (UMAP), to 265 FFPE (**a**) and 568 FFPE (**b**) sWGS samples, using the top ten key indel signature features as shown in Fig. 4f. The MSI status of ‘0’ refers to non-MSI/MMR-proficient (MMRp) samples (blue dots), and the MMR status of ‘1’ refers to MSI/MMRd samples (red dots). Different dot shapes indicate different data sources, as shown in the figure legend.

**Supplementary Fig. 21.**
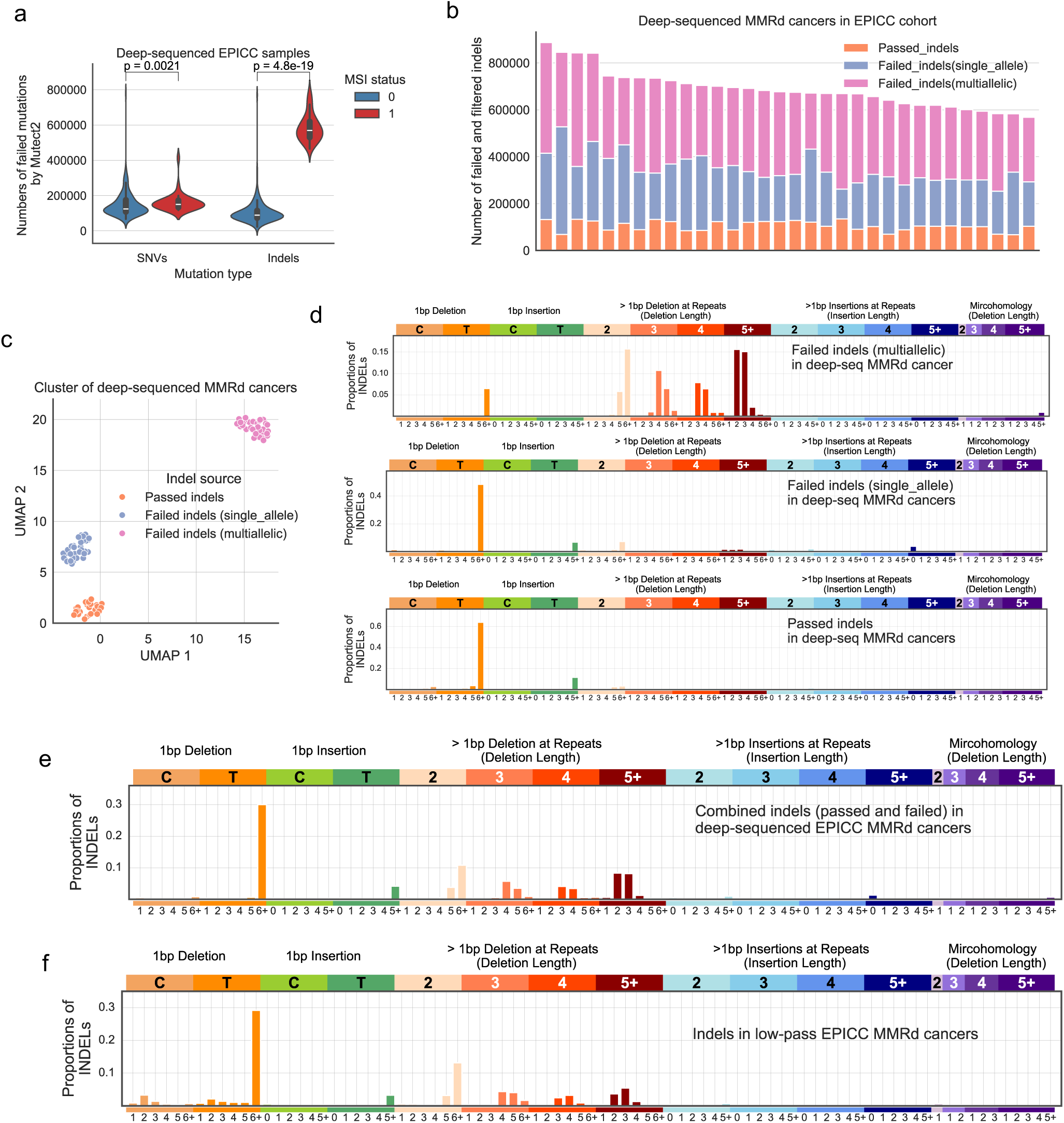
Longer deletions can be subject to bias filtering by the mutation caller in deep-sequenced MMRd samples. (**a**) Violin plot comparing the failed mutation count between MSI and non-MSI samples from the EPICC cohort. The filtered mutations were those without a ‘PASS’ tag in raw mutation list called by Mutect2. The width of the violin plot reflects the frequency of data points in each region, and the overlaid box plot in the centre indicates the interquartile range, with the white dot representing the median. (**b**) Stacked bar chart illustrating the distribution of raw indel mutations identified in 32 deep-sequenced MSI samples. Passed indels: orange; failed indel mutations with a single allele: blue; failed indels with multiple alleles: purple. (**c**) Clustering analysis of 32 EPICC MSI samples using different groups of indel mutations. (**d**) Indel mutation profiles across different groups of indel mutations in the 32 deep-sequenced MSI EPICC samples. (**e**) Combined indel mutation profile of deep-sequenced MSI samples. (**f**) Mean indel mutation profile of sWGS EPICC MMRd samples. It is provided here for reference to panel (**e**), which is also included as top panel plot in Supplementary Fig. 18b.

**Supplementary Fig. 22.**
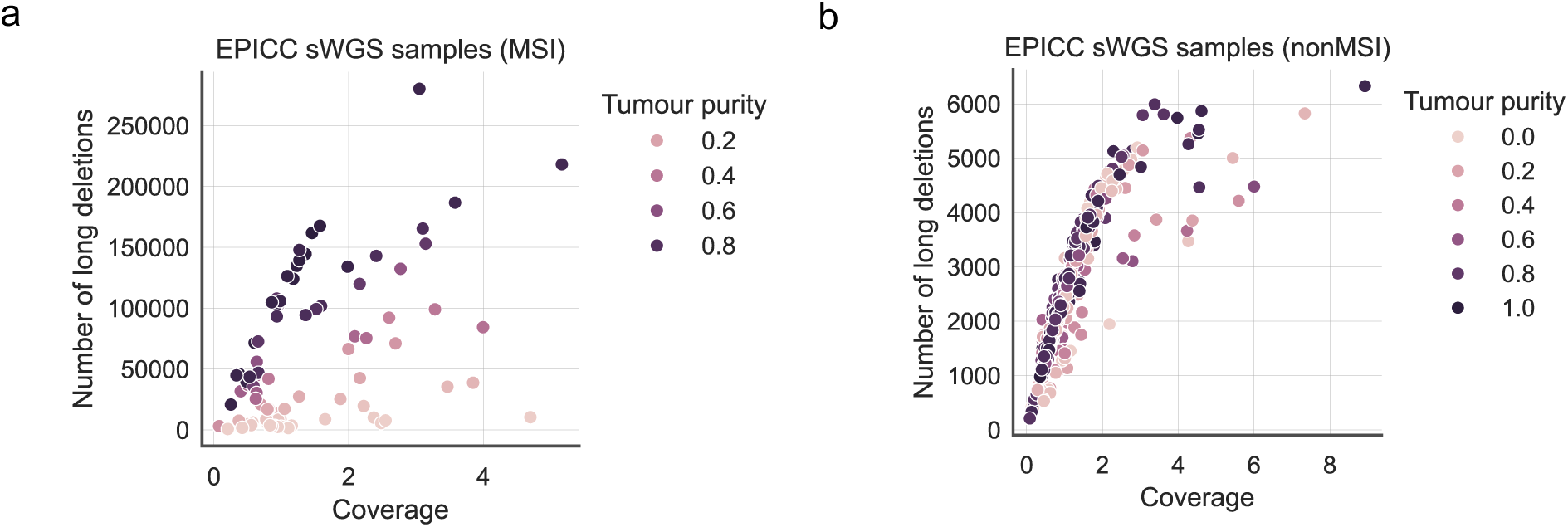
Correlation between long-deletion count and the sequencing coverage in FF MSI (a) and non-MSI (b) sWGS samples. The long-deletion count is calculated as the sum of mutation counts in key long-deletion signature channels. We show the observed long-deletion count on *Y*-axis and the sequencing coverage on X-axis. We coloured each dot using the sample tumour purity.

**Supplementary Fig. 23.**
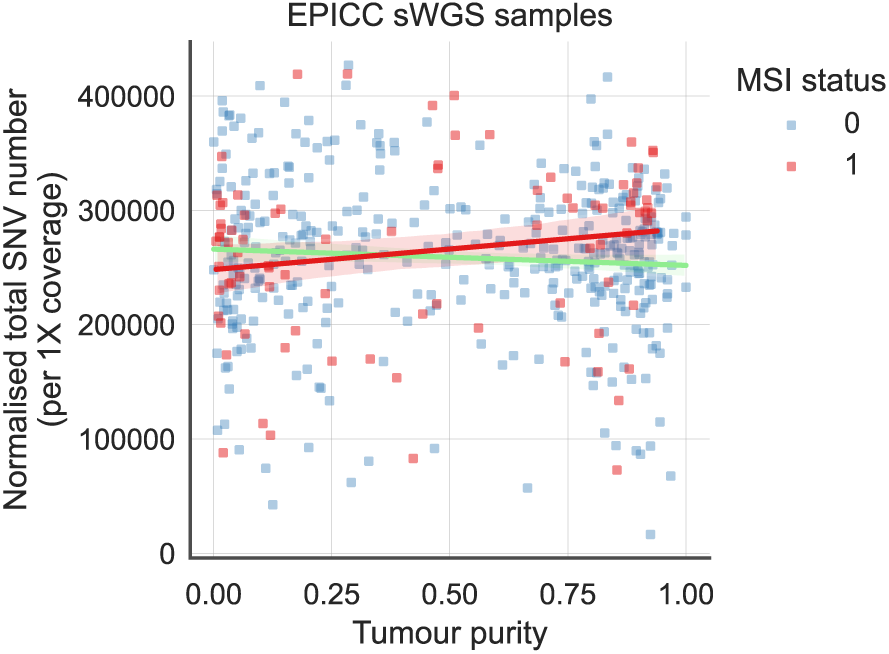
Relationship between the normalised SNV count and tumour purity in FF MSI and non-MSI sWGS samples. The total SNV is normalised by dividing the sequencing coverage. The red and green lines show linear regression fits for MSI and non-MSI samples.

**Supplementary Fig. 24.**
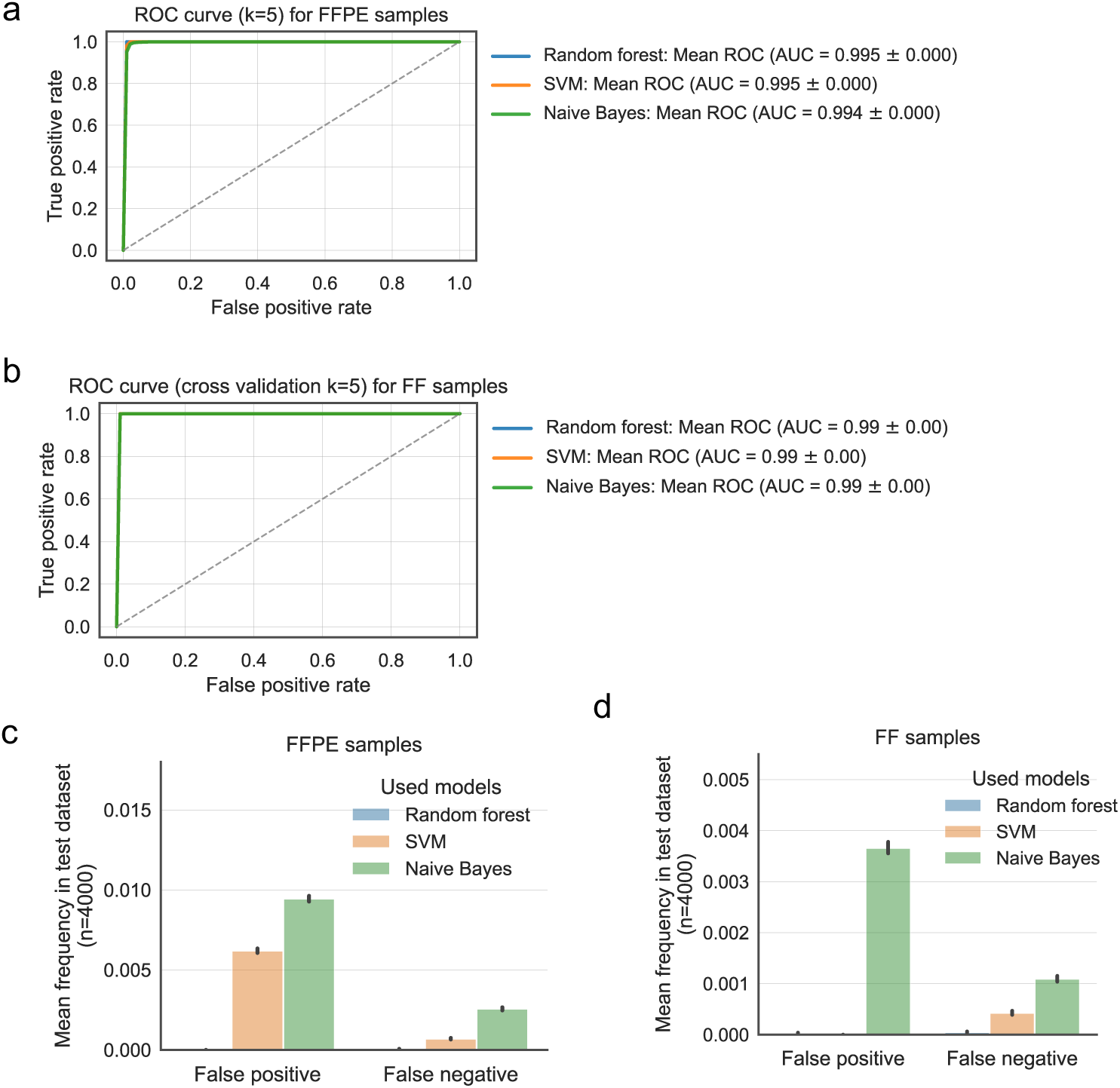
Model performance of three classifiers on synthetic sWGS samples. **(a-b)** ROC curves for MMRd classification by three different classifiers in FFPE (**a**) and FF (**b**) sWGS samples. The classifiers were trained on 80% of the simulated samples and tested on the remaining 20% using 5-fold cross-validation. The AUC values and standard deviations are shown in each figure legend. (**c-d**) False prediction rates of the classifiers in FFPE (**c**) and FF (d) sWGS samples. We repeated the 5-fold cross-validation 50 times with different random initiations for both FFPE and FF samples, collecting false positive and false negative rates on the test samples for each setting. Error bars represent the standard deviations.

**Supplementary Fig. 25.**
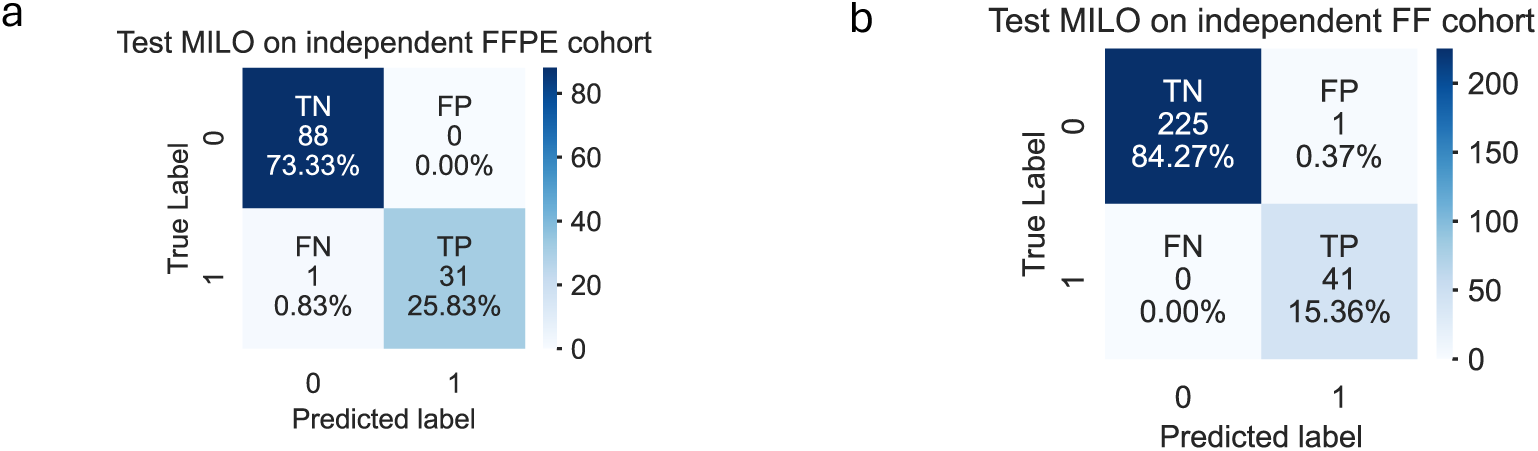
Confusion matrix of random forest classifier on the third independent FFPE (a) and FF (b) sWGS cohort. The matrices display true negatives (TN), true positives (TP), false positives (FP), and false negatives (FN).

**Supplementary Fig. 26.**
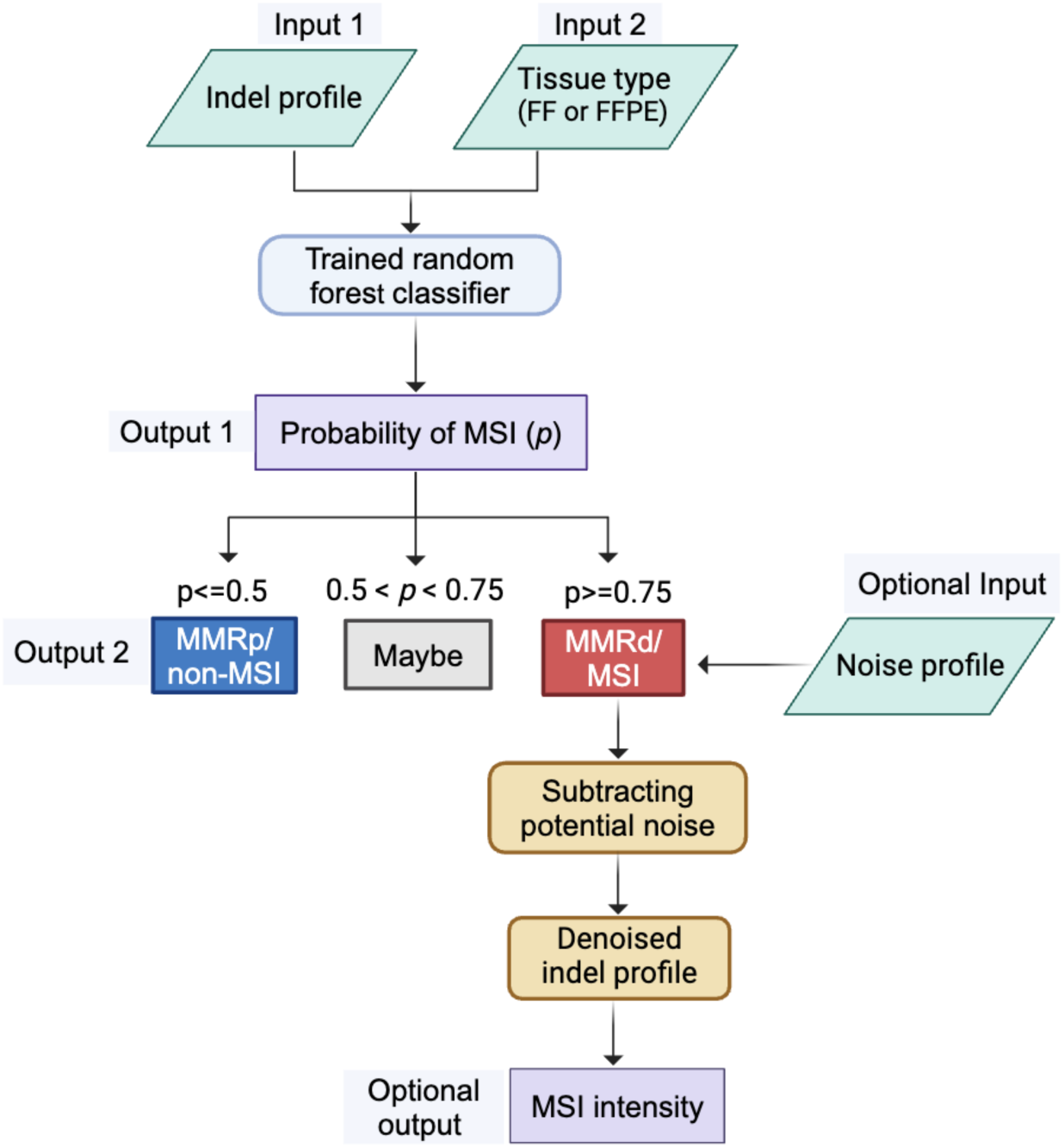
Workflow of MILO. MILO employs two random forest classifiers trained on both simulated and real sWGS data from FFPE and FF samples. The workflow begins with two key inputs: the ‘observed indel profile’ and the tissue type (’FFPE’ or ‘FF’), which determines the appropriate classifier. The classifier calculates the MSI probability and categorises the sample into one of three groups: ‘MSI,’ ‘Maybe,’ or ‘non-MSI,’ based on the MSI probability as shown in the workflow. Optionally, MILO can perform noise subtraction using either the default or a customised noise profile, and then reports the MSI intensity based on the noise-adjusted profile. A detailed user manual is available in https://github.com/QingliGuo/MILO.

